# Control of nuclear size by osmotic forces in *Schizosaccharomyces pombe*

**DOI:** 10.1101/2021.12.05.471221

**Authors:** Joёl Lemière, Paula Real-Calderon, Liam J. Holt, Thomas G. Fai, Fred Chang

## Abstract

The size of the nucleus scales robustly with cell size so that the nuclear-to-cell volume ratio (N/C ratio) is maintained during cell growth in many cell types. The mechanism responsible for this scaling remains mysterious. Previous studies have established that the N/C ratio is not determined by DNA amount but is instead influenced by factors such as nuclear envelope mechanics and nuclear transport. Here, we developed a quantitative model for nuclear size control based upon colloid osmotic pressure and tested key predictions in the fission yeast *Schizosaccharomyces pombe*. This model posits that the N/C ratio is determined by the numbers of macromolecules in the nucleoplasm and cytoplasm. Osmotic shift experiments showed that the fission yeast nucleus behaves as an ideal osmometer whose volume is primarily dictated by osmotic forces. Inhibition of nuclear export caused accumulation of macromolecules and an increase in crowding in the nucleoplasm, leading to nuclear swelling. We further demonstrated that the N/C ratio is maintained by a homeostasis mechanism based upon synthesis of macromolecules during growth. These studies demonstrate the functions of colloid osmotic pressure in intracellular organization and size control.

## Introduction

It has been known for more than a century that the size of the nucleus scales with cell size. Since the initial observation in plants (Strasburger, 1893) the scaling of nuclear and cell volume has been documented across the eukaryotic domain (Conklin, 1912; Gregory, 2005; Moore et al., 2019; Neumann and Nurse, 2007). More recently, scaling was even observed for nucleoids in prokaryotes (Gray et al., 2019). In multicellular organisms, the nuclear-to-cell volume (N/C) ratio varies among cell types, but this ratio is maintained as a constant within a given cell type (Conklin, 1912; Hertwig, 1903). During cell growth, the N/C ratio is also maintained through much of the cell cycle (Jorgensen et al., 2007; Neumann and Nurse, 2007; Willis et al., 2016), as the nucleus grows in volume at the same rate as the cell. Abnormal N/C ratios are a hallmark of diseases such as certain cancers and are sometimes used as diagnostic criteria (Foraker, 1954; Slater et al., 2005; Webster et al., 2009; Zink et al., 2004). The N/C ratio may play an important role in regulatory mechanisms, for instance, in the mid-blastula transition in embryonic development (Amodeo et al., 2015; Jevtić and Levy, 2015). However, despite the universal and fundamental nature of this cellular property, the mechanistic basis for nuclear size scaling remains poorly understood.

Although there is a correlation between nuclear size and amount of DNA, it is weak and therefore unlikely that DNA itself is the responsible scaling factor. DNA is only a minor component in the nucleus by volume; it has been estimated to occupy <1% of the nuclear volume and is many times less abundant in the nucleus than RNA (Milo and Phillips, 2015). Nuclear size does increase with increased ploidy in a given cell type, but generally this increase is accompanied by a similar increase in cell size (Cavalier-Smith, 2005; Gregory, 2005; Gregory and Mable, 2005; Jorgensen et al., 2007; Robinson et al., 2018). During the cell cycle, nuclear size continues to grow in the G2 phase even when DNA content is no longer increasing (Jorgensen et al., 2007; Neumann and Nurse, 2007). Further, through manipulating genome content in fission yeast, it has been shown that cells with DNA content ranging from 2N to 32N have a similar N/C ratio (Neumann and Nurse, 2007). Thus, DNA is unlikely to be the rate-limiting structural component that determines nuclear size.

Nuclear size and shape are dictated both by nuclear volume and surface area. It is clear however that nuclear volume and surface area can be uncoupled and are regulated independently. For instance, arrest of budding yeast cells in mitosis can lead to continued growth of the nuclear envelope without growth in nuclear volume, leading to misshapen nuclei and formation of nuclear envelope protrusions (Webster et al., 2009). Growth of the nuclear envelope may occur through the transfer of membranes from the endoplasmic reticulum or by lipid assembly at the nuclear envelope (Blank et al., 2017; Hirano et al., 2020; Kim et al., 2007). Studies have shown however that nuclear volume, not surface area, is the relevant geometric parameter that is maintained for the N/C ratio (Cantwell and Nurse, 2019a; Neumann and Nurse, 2007; Walters et al., 2019).

Efforts to define molecular-based control mechanisms have been largely unsuccessful. Genome-wide screens in fission yeast have demonstrate that the vast majority of mutants exhibit normal N/C ratios, ruling out many possible cellular processes and molecular pathways (Cantwell and Nurse, 2019b; Kume et al., 2017). For instance, the N/C ratio is independent of cell size, shape, and number of nuclei (Neumann and Nurse, 2007). Screens have so far identified only a small number of genes that impact the N/C ratio, mostly related to nuclear transport or lipid synthesis (Cantwell and Nurse, 2019b; Kume et al., 2017). In vertebrate cell systems, lamins and chromatin factors have been implicated in the control of nuclear size and shape (Edens et al., 2017; Levy and Heald, 2010; Muchir et al., 2004). For example, depletion of lamin in *Xenopus* eggs extract resulted in a reduction of nuclear size and formation of abnormal nuclear shapes (Newport et al., 1990). However, as yeast lack lamins, it is unlikely that the nuclear lamins themselves represent a universal mechanism for nuclear size control.

Another potential factor in nuclear size control is osmotic pressure. Instead of a rigid structure, the nucleus may be regarded as a structure similar to a balloon whose size is dependent on the balance of pressures and membrane tension. The rounded shape of the typical nucleus suggests there may be slightly higher osmotic pressure in the nucleoplasm compared to the cytoplasm, which is balanced by the nuclear membrane tension. These pressures likely arise from macromolecular crowding forces termed “colloid osmotic pressure,” which are produced by the distinct sets of macromolecules in the nucleus and cytoplasm (Mitchison, 2019). The osmotic nature of the nucleus has been shown in various ways. Treatment of cells with osmotic shocks causes both the cell and nucleus to swell and shrink (Churney, 1942). Classic experiments demonstrated that injection of crowding agents such as polyethylene glycol into the cytoplasm cause shrinkage of the nucleus (Harding and Feldherr, 1958, 1959). Isolated nuclei are also responsive to osmotic shifts but the osmotic behavior depends on the molecular size of the osmolytes such that only macromolecules larger than 30 kDa will affect their volumes (Finan et al., 2009). In general, a rigorous quantitative assessment of the osmotic model for nuclear size control is lacking.

Here, we developed a quantitative model for nuclear size control based upon osmotic forces, using a combination of theoretical modeling and quantitative experiments. We used fission yeast as a tractable model in which cellular and nuclear volumes can be accurately measured. We propose a theoretical framework that represents the nucleus and cell as a system of nested osmometers. We show that nuclei in fission yeast behave as ideal osmometers, which allows for the direct study of the effects of osmotic pressure on nuclear volume and its responses to changes in macromolecular crowding. This osmotic model suggests a mechanism for maintenance of the N/C ratio during cell growth, as well for homeostasis behavior that corrects an aberrant N/C ratio over time. Together, these studies provide critical quantitative support for an osmotic-based mechanism for nuclear size control.

## Results

### Model of the nucleus and a cell as two nested osmometers

We developed a quantitative model of nuclear and cell size control based on the physical mechanism of osmosis. The nucleus and the cell are represented as a system of nested osmometers, whose volumes are determined by osmotic pressure differences, membrane tensions, and non-osmotic volumes (Figure 1A). The cell is inflated by turgor pressure, which is defined as the osmotic pressure difference across the plasma membrane (C^out^, C^Cy^) balanced by the elastic wall surrounding the cell. This turgor pressure is produced largely from small molecules, such as ions and metabolites, attracting water into the cell through osmosis. The nuclear envelope is a semi-permeable membrane that allows water and ions and other small molecules (with a Stoke radius below <2.5 nm (Mohr et al., 2009)), but remains relatively impermeable to large proteins, macromolecular complexes, DNA and RNA, with the exception of specific nuclear transport mechanisms through nuclear pores. Certain macromolecules produce colloid osmotic pressures, in part by organizing a shell of water and ions around them (Mitchison, 2019). Only molecules that are too large to freely diffuse across the nuclear envelope produce colloid osmotic pressure (*π*^cy^, *π*^N^), whereas small molecules that are freely diffusing between the cytoplasm and nucleus do not. In addition, there are also non-osmotically active volumes in the cytoplasm and nucleoplasm (b^Cy^, b^N^), which represents the dry volume taken up by cellular components; this term also describes the degree of macromolecular crowding.

**Figure 1.**
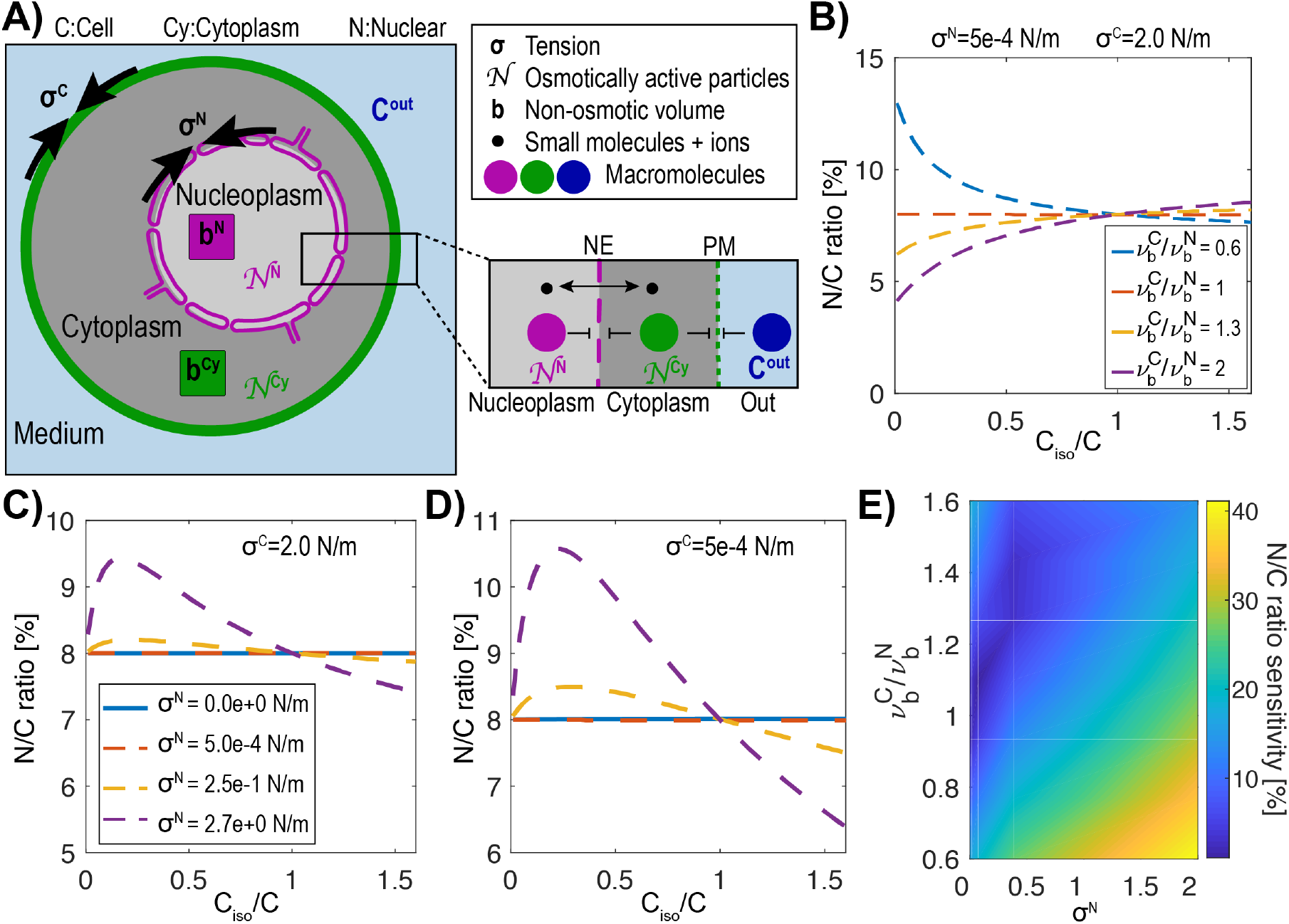
Model of the nucleus and the cell as “a vesicle within a vesicle”, osmotically challenged. (**A)** Schematic of the model and parameters used in the mathematical model: membrane tension σ, non-osmotic volume b, number (N) of macromolecules that cannot freely cross either the cell or nuclear membranes, concentration of the buffer C^out^. (**B)** Theoretical prediction of the effect of a change in the normalized inverse external concentration C_iso_/C on the N/C ratio for various ratios of normalized cellular (*v_c_*) and nuclear (*v*_N_) non-osmotic volume values keeping the cell and nuclear membrane tensions (σ^C^, σ^N^) constant. (**C)** Predictions of osmotic shifts on the N/C ratio for various nuclear membrane tensions (σ^N^), keeping a high cell tension (σ^C^) constant. (**D)** Same as (C) keeping a low membrane tension. (**E)** Phase diagram of the N/C ratio sensitivity to osmotic shocks defined as [max(N/C ratio)-min(N/C ratio)] / (N/C ratio_isotonic_) for various ratios of non-osmotic volumes and nuclear membrane tension.

We postulated that the nucleus is inflated by colloid osmotic pressure differences between the nucleoplasm and cytoplasm; such differences are estimated to be orders of magnitude smaller than turgor pressure (kPa versus MPa in yeast). The expansion of the cell and nuclear membranes can also be restricted by membrane tension (σ). At the cell surface, σ^C^ includes plasma membrane tension as well as other mechanically relevant features such as the cell wall or cortex. Similarly, for the nuclear envelope, σ^N^ includes the lipid bilayer tension and potentially the mechanical properties of the lamina and chromatin anchored to the membrane (Schreiner et al., 2015). Other factors are known to alter membrane tension, such as membrane reservoirs allowing increases in membrane surface area while keeping membrane tension low. Examples of possible reservoirs include invaginations of the plasma membrane such as eisosomes (Lemière et al., 2021) and caveolae (Sinha et al., 2011) and the endoplasmic reticulum for the nuclear envelope. We used established osmotic theory, including Boyle Van’t Hoff’s relationship and Laplace’s Law (Laplace, 1805), to analyze the behavior of osmometers in our model. We treated the cell and nucleus as two spherical nested osmometers having respective membrane tensions σ^C^ and σ^N^ and interpreted Van ‘t Hoff’s Law in terms of the concentrations of osmotically active particles in the cytoplasm (C^Cy^), nucleoplasm (C^N^), and extracellular space (C^out^). We described the steady state solutions in which colloid osmotic pressures in the cytoplasm and nucleus are in balance with their respective membrane tension, which results in the coupled equations:

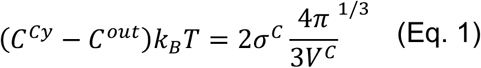

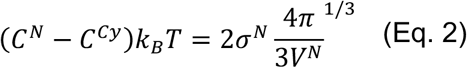

Solving this system of equations for the unknown cell volume (*V^C^*) and nuclear volume (*V^N^*) yields a unique steady-state value for the N/C ratio (Appendix A). In some special cases, the N/C ratio may be written explicitly in terms of the parameters, as we show later on. However, in general the equations are solved numerically. Note that ions do not contribute to the osmotic balance in Eq. 2. as they are permeable to the nuclear envelope.

Using this model, we evaluated what key parameters affect the N/C ratio. To do this, we solved this system of equations (Appendix B Eqs. 14 and 15) for different sets of parameters to find the resulting N/C ratio. One prediction of this model is that if 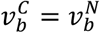 which represent the normalized non-osmotic volume of the cell and the nucleus (defined as 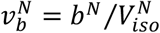 and 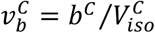) then the N/C ratio remains constant under osmotic shifts (Figure 1B). Conversely, whenever 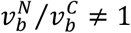, the model predicts that the N/C ratio will vary with osmotic shocks (Figure 1B). Another key prediction is that if σ^N^ = 0 N/m, the N/C ratio remains constant upon osmotic shifts (Appendix B Eq. 16). In Figures 1C and D we plotted the effects of nuclear membrane tension σ^N^ (from 0 to 2.7 N/m) on the N/C ratio upon osmotic shifts. The results also reveal that the N/C ratio is relatively insensitive to osmotic shocks for small values of σ^N^ independently of σ^C^ (σ^N^ = 0.5 mN/m, Figures 1C, 1D, Appendix B and C). Figure 1E summarizes these findings on the effects of varying both σ^N^/ σ^C^ and 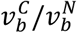.

We considered a special case where nuclear membrane tension σ^N^ = 0 N/m, and the normalized non-osmotic volume in the nucleus and cytoplasm are balanced, with 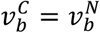. As explained in Appendix D, in this case the N/C ratio is set simply by the ratio of the numbers of osmotically active molecules in the nucleoplasm and cell:

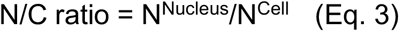

In the sections below, we tested and further developed this osmotic-based model with experiments with fission yeast to measure key parameters and test model predictions.

### The *S. pombe* nucleus behaves as an ideal osmometer

To quantify the osmotic forces that control cell and nuclear size, we experimentally determined the volume responses of fission yeast cells and their nuclei to osmotic shifts in their media. To visualize the cell and nucleus, we imaged fission yeast cells expressing a nuclear membrane marker (Ish1-GFP, (Expósito-Serrano et al., 2020)) and a plasma membrane marker (mCherry-Psy1,(Kashiwazaki et al., 2011))(Figure 2A, B). We placed live cells in flow chambers and treated them with media containing various concentrations of sorbitol, an osmotic agent (see Methods). Nuclear and cell volumes were measured using a semi-automated 3D segmentation approach (Methods; Figure S1A). As cells adapt to hyperosmotic shocks by gradually increasing glycerol production to recover their volume (Chen et al., 2003), we minimized these adaptation effects by taking measurements acutely upon shocks (<1 min) and by using a *gpd1*Δ mutant background that is delayed in this response (Hohmann, 2002; Minc et al., 2009) (Figures S1A-C). To analyze volume responses, we used Boyle Van’t Hoff (BVH) plots in which the normalized inverse volumes are plotted as a function of normalized inverse concentration in medium (Figure 2C). Ideal osmometers are characterized by linear responses following BVH’s Law (Nobel, 1969), showing that their volume is determined primarily by the osmotic environment with negligible effects of surface tension (Figure 2C; dotted line). In contrast, in cases with significant membrane tension, the plots exhibit non-linear responses (Figure 2C; green line). Further, the intersection of the BVH plot at the Y-axis provides a measure of the normalized non-osmotic volume (*v_b_*; Figure 2C).

**Figure 2.**
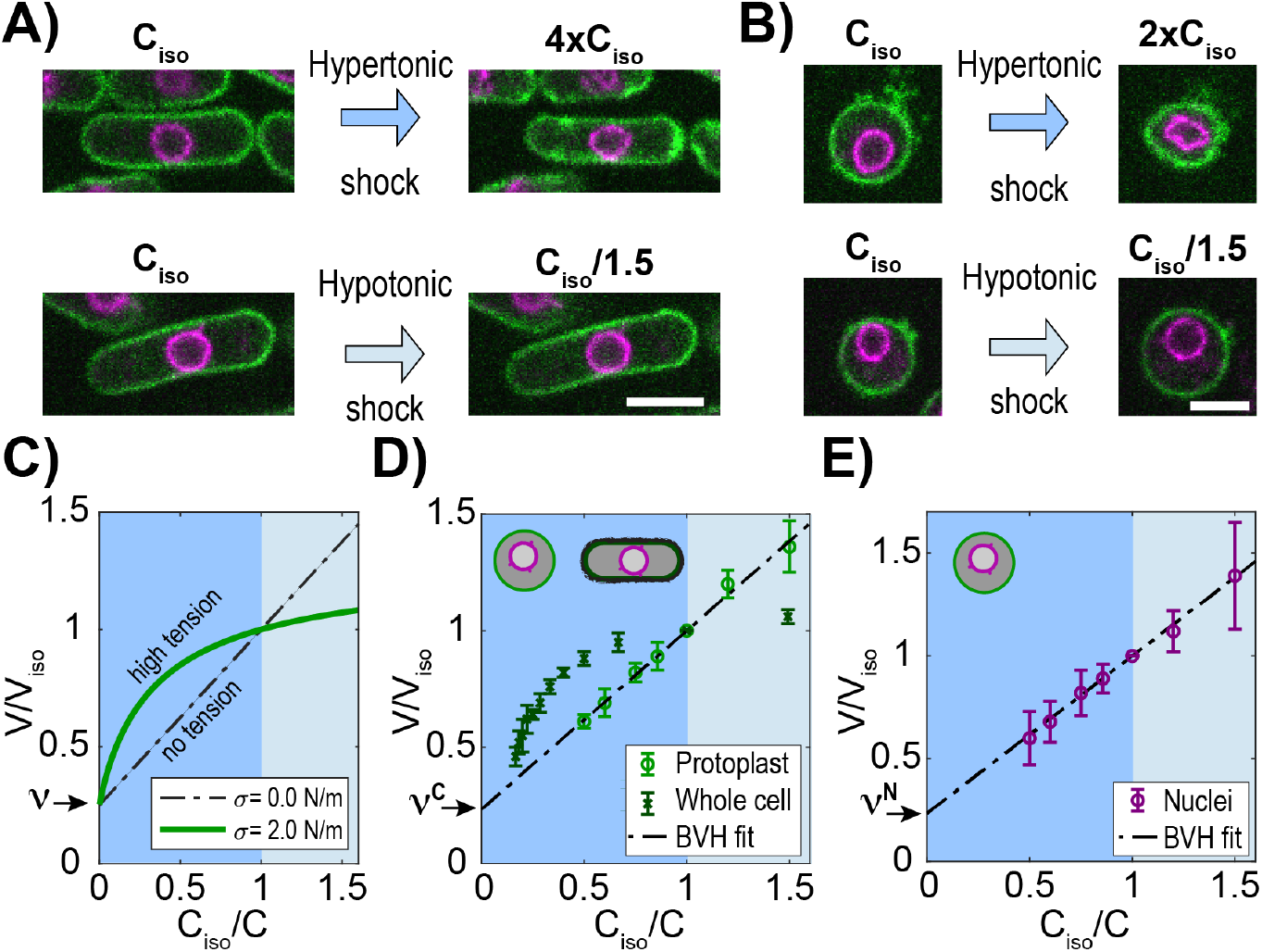
The fission yeast nucleus behaves as an ideal osmometer. **(A)** Images of cells expressing a plasma membrane marker mCherry-Psy1 (green) and a nuclear envelope marker Ish1-GFP (purple). Individual cells in isotonic medium (C_iso_) were shifted to hypertonic or hypotonic medium and imaged for 3D volume measurements (Methods). **(B)** Images of individual protoplasts in response to hypertonic and hypotonic shifts. See also Figure S2. Scale bar = 5 μm. **(C-E)** BVH plots of the effects of osmotic shifts on the volume of the cell and nucleus. **(C)** Theoretical predictions of effects of osmotic concentration in the medium (C_iso_/C) on the volume of a cell or nucleus with zero (black) or large (green) membrane tension (σ). Dashed line (black) depicts the behavior of an ideal osmometer in which there is no effect of membrane tension. **(D)** Effect of osmotic shifts on the relative volumes (V/V_iso_, mean ± STD) of whole fission yeast cells (N=707, 3 biological replicates) and protoplasts (N=441, from at least 5 biological replicates). **(E)** Effect of osmotic shifts on relative nuclear volume (V/V_iso_, mean ± STD) in protoplasts (N=441, from at least 5 biological replicates). Note that the response of nuclei fits to the predicted behavior of an ideal osmometer.

First, we analyzed the effect of osmotic shifts on cellular volume. Hyperosmotic shifts of various sorbitol concentrations caused sizable decrease (up to ~54%) in volume of cells, as previously noted (Atilgan et al., 2015) (Figures S1B-C). The BVH plot showed that the response was non-linear, indicative of a non-ideal osmometer behavior (Figure 2D). The relationships were non-linear for both hyper- and hypotonic responses, consistent with the actions of the elastic cell wall that exerts compressive forces on the cell body and resists large expansion of volume (Atilgan et al., 2015; Davì and Minc, 2015; Schaber and Klipp, 2008).

We therefore conducted osmotic shift experiments on protoplasts, which are yeast cells in which the cell walls has been enzymatically removed (Flor-Parra et al., 2014a; Lemière et al., 2021) (Figure 2B). To maintain viability of protoplasts, sorbitol was added to the medium as osmotic support to substitute for the role of the cell wall and to prevent lysing. We determined the isotonic conditions for these protoplasts to be YE medium supplemented with 0.4 M sorbitol (hereafter called YE + 0.4 M), as they had similar cytoplasmic properties as walled cells in YE + 0 M sorbitol. At this concentration of sorbitol, an asynchronous population of protoplasts exhibited similar average volumes as those of walled cells in YE + 0 M sorbitol, and similar cytoplasmic concentrations as assessed by fluorescence intensity of a cytoplasmic marker E2-mCrimson (Figures S2A-B) (Knapp et al., 2019). For osmotic shift experiments, we prepared protoplasts in this isotonic condition of YE + 0.4 M sorbitol (C_iso_), and then shifted them into medium containing a range of sorbitol concentrations below and above the isotonic condition (0.2 to 1.0 M). These methods allowed for quantitative probing of osmotic effects over a remarkable ~3-fold range of volume; notably, protoplasts were able to swell up to 40% in volume or shrink 40% without bursting.

The BVH plot of protoplast responses showed a linear behavior through this range of sorbitol concentrations (Figure 2E), indicative of an ideal osmometer. As the number of osmolytes is directly related to cell volume in osmotic shift experiments for ideal osmometers, this allowed us to estimate *S. pombe* solute concentration at ~30×10^7^ solutes/μm^3^, which represents an osmolarity of 500±45 mOsmol (Figures S2C-G). The BVH plot also showed the normalized non-osmotic volume 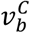 to be 25%, similar to what has been previously reported for fission yeast cells (Atilgan et al., 2015) and other organisms (Dill et al., 2011; Ellis, 2001).

Having found that protoplasts behave as ideal osmometers, we then measured their responses in nuclear volume. In hyperosmotic shifts, nuclei in both whole cells and protoplasts shrank into an abnormal involuted shape, suggesting a loss in volume but not surface area (Figure 2A, B), similar to what has been observed in mammalian cells (Kim et al., 2016). In hypoosmotic shifts with protoplasts, the nuclei expanded in a spherical shape to increase 40% in volume and 26% in surface area (Figure 2B, E). Strikingly, this large expansion of the nuclear envelope occurred in < 1 min, suggesting that the nuclear envelope can rapidly draw upon membrane stores, potentially from the endoplasmic reticulum (Kume et al., 2019; Roubinet et al., 2021). BVH plots showed that the volume of nuclei in protoplasts followed a linear behavior in osmotic shifts over a 3-fold range of volumes (Figure 2E). Importantly, this linear response showed that the nucleus behaved as an ideal osmometer. This finding implies that tension of the nuclear envelope is negligible: σ^N^ ≈ 0 N/m; the nuclear envelope does not exert tension that alters the volume response to osmotic forces, so that nuclear volume is directly responsive to its osmotic environment. The BVH plot also revealed that the normalized non-osmotic volumes in the nucleus 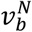 and cytoplasm 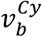 are similar (25% in nucleus; 25% in cytoplasm) (Figure 2D, E).

Thus, these experimental findings show that the protoplast and the nucleus approximate two nested spherical ideal osmometers as described in our theoretical model. As the physical properties of the nucleus are unlikely to change in the short amount of time needed to remove the cell wall, these results imply that the nucleus in whole cells (those with intact cell walls) are also ideal osmometers. Taken together, these findings indicate that fission yeast cells may be represented by the simplest version of the model where σ^N^ ≈ 0 N/m with matching 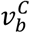 and 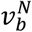, so that the N/C ratio is determined directly by the ratio of osmotically active molecules in the nucleoplasm to cell.

### The N/C ratio does not depend on the presence of the cell wall

According to the model, the N/C ratio should be independent of the external tension (σ^C^) and the outside concentration (C^out^) (Figure 1). To reduce σ^C^ we examined the effects of removing the cell wall. The fission yeast cell wall has an elastic surface modulus of σ^C^~10 to 20 N/m which resists 1.5 MPa of turgor pressure (Atilgan et al., 2015; Minc et al., 2009). Upon removal of the cell wall, protoplasts are maintained in medium with sorbitol and have a five-orders-of-magnitude decrease in σ^C^, with a membrane tension of ~4.5×10^−4^ N/m (Lemière et al., 2021). We tracked individual cells during cell wall digestion as they were converted to protoplasts (Figure 3A, right panel). There was no significant change in the N/C ratio before and after cell wall removal (Figure 3A, left panel). We noted that N/C ratios were slightly elevated in our initial population measurements, but this effect was due to loss of cellular material during the process of protoplasting (Figure S3A). Thus, as predicted by our model, the N/C ratio is independent of the outer tension of the system (σ^C^).

**Figure 3.**
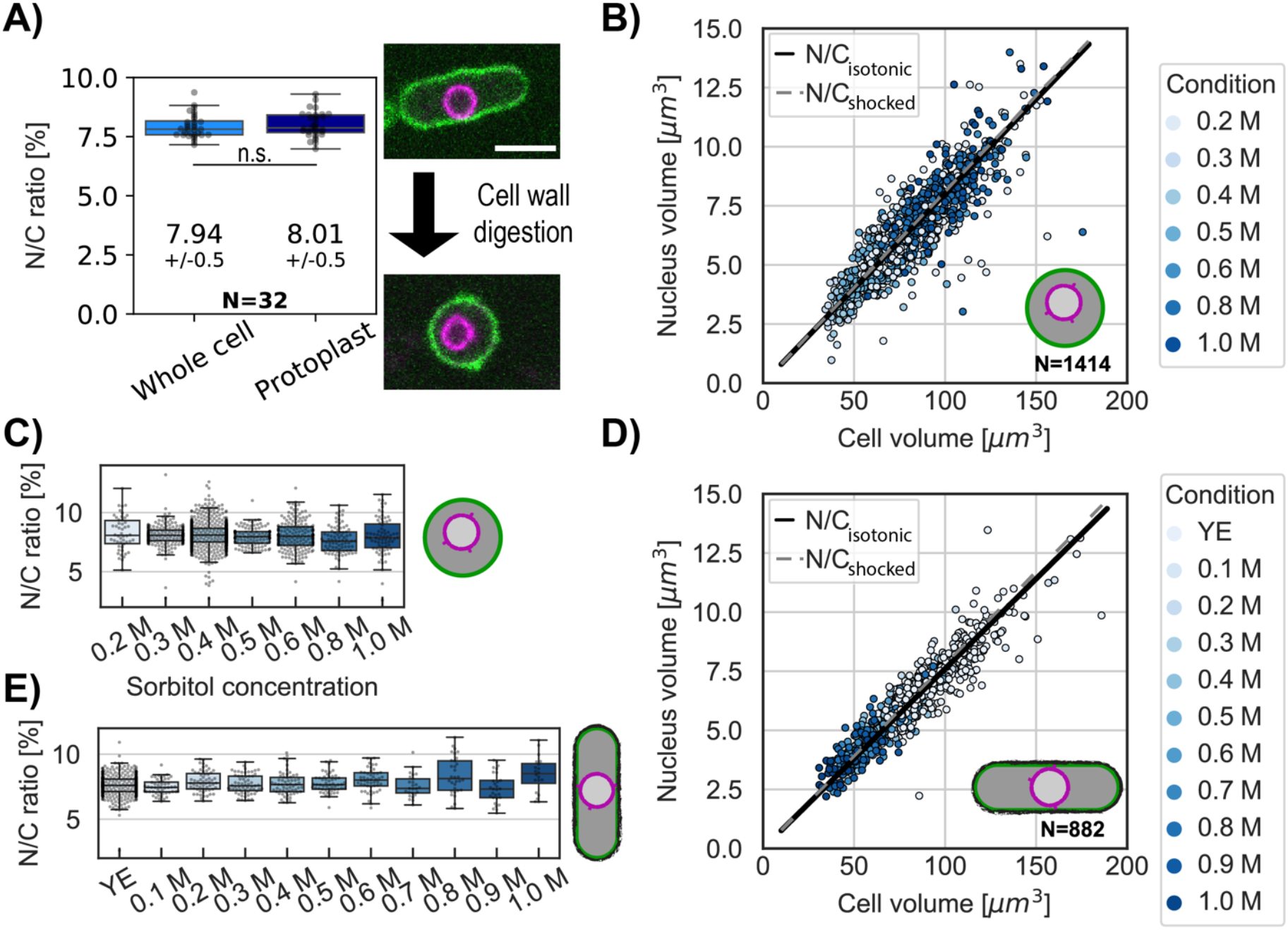
The N/C ratio is maintained in osmotic shifts and upon cell-wall removal. **(A)** The N/C ratio of the same cells before and after cell wall digestion (mean ±STD) reveals no statistical differences (paired t test, p=0.36). Right panel, overlay of the plasma membrane (green) and nuclear membrane (purple) of the same cell middle plane before and after cell-wall digestion. For all box and whiskers plots, the horizontal line indicates the median, the box indicates the interquartile range of the data set (IQR) while the whiskers show the rest of the distribution within 1.5*IQR except for points that are defined as outliers. Scale bar = 5 μm. From 6 biological replicates. (**B)** Scatter plot of cell size and nuclear size for protoplasts under isotonic conditions (YE + 0.4 M sorbitol) and immediately following osmotic shocks. Black and dashed gray lines, measured N/C ratio of cells under isotonic conditions and osmotically shocked, respectively. From at least 2 biological replicates. **(C)** Protoplasts with the individual N/C ratio per osmotic condition described in (B). **(D)** Same as (B) for whole cells in YE under isotonic condition. Black line, measured N/C ratio of cells in isotonic conditions. From 2 biological replicates. **(E)** Same as (C) for whole cells with the individual N/C ratio per osmotic condition described in (D). See also Figure S3.

### The N/C ratio is maintained under osmotic shifts in protoplasts and whole cells

An important prediction of the osmotic model is that the N/C ratio should not change upon osmotic shifts. To test this prediction, we subjected protoplasts to a range of hypo and hyper shocks and measured nuclear and cellular volumes. We varied sorbitol concentrations from 0.2-1.0 M with isotonic conditions defined as 0.4M sorbitol (Figure S2A, S2B). A plot of nuclear versus cellular volumes showed scaling was robustly maintained throughout the range of osmotic conditions (Figures 3B, S3B-C). This was also shown by measurements of the N/C ratios at each sorbitol concentration (Figure 3C). Similar experiments in whole cells showed that the distribution of the N/C ratio under osmotic shock (0.1M to 1.0 M) also coincided with the distribution of the same population of whole cells in isotonic condition (Figures 3D-E, S3D-E). These results demonstrated that the N/C ratio does not change with the osmotic concentration of the media, confirming the predictions of the model that the sizes of the cell and nucleus are both regulated by osmotic pressures.

### Nanorheology reveals that physical properties of the cytoplasm and nucleoplasm are comparable under osmotic shocks

A key test of the model is to experimentally measure the relevant macromolecular concentrations and intracellular colloid osmotic pressures in both the cytoplasm and nucleoplasm. Recent advances have facilitated measurements of these parameters (Mitchison, 2019). We used forty nanometer-sized genetically encoded multimeric nanoparticles (GEMs) labeled with mSapphire fluorescent protein as nanorheological probes to quantitatively measure macromolecular crowding through analyses of their diffusive motions (Delarue et al., 2018; Knapp et al., 2019; Molines et al., 2020) (Figure 4A, green). We used two versions of the GEMs: cytGEM and nucGEM, to measure crowding in the cytoplasm and nucleoplasm, respectively. The nucGEM protein is a version of GEMs that contains a nuclear localization signal (NLS) (Szoradi et al., 2021); the NLS-GEM monomer is thought to be transported into the nucleus and retained once it assembles into the nucleus with the NLS embedded inside the spherical particle. Cells expressing this NLS-GEMs fusion exhibited motile fluorescent particles in the nucleus (Figure 4A). Projections of images over time showed that the nuclear GEMs were excluded from the nucleolus, so that nuclear GEMs primarily probe the properties of non-nucleolar portion of the nucleus (Figure 4A, purple).

**Figure 4.**
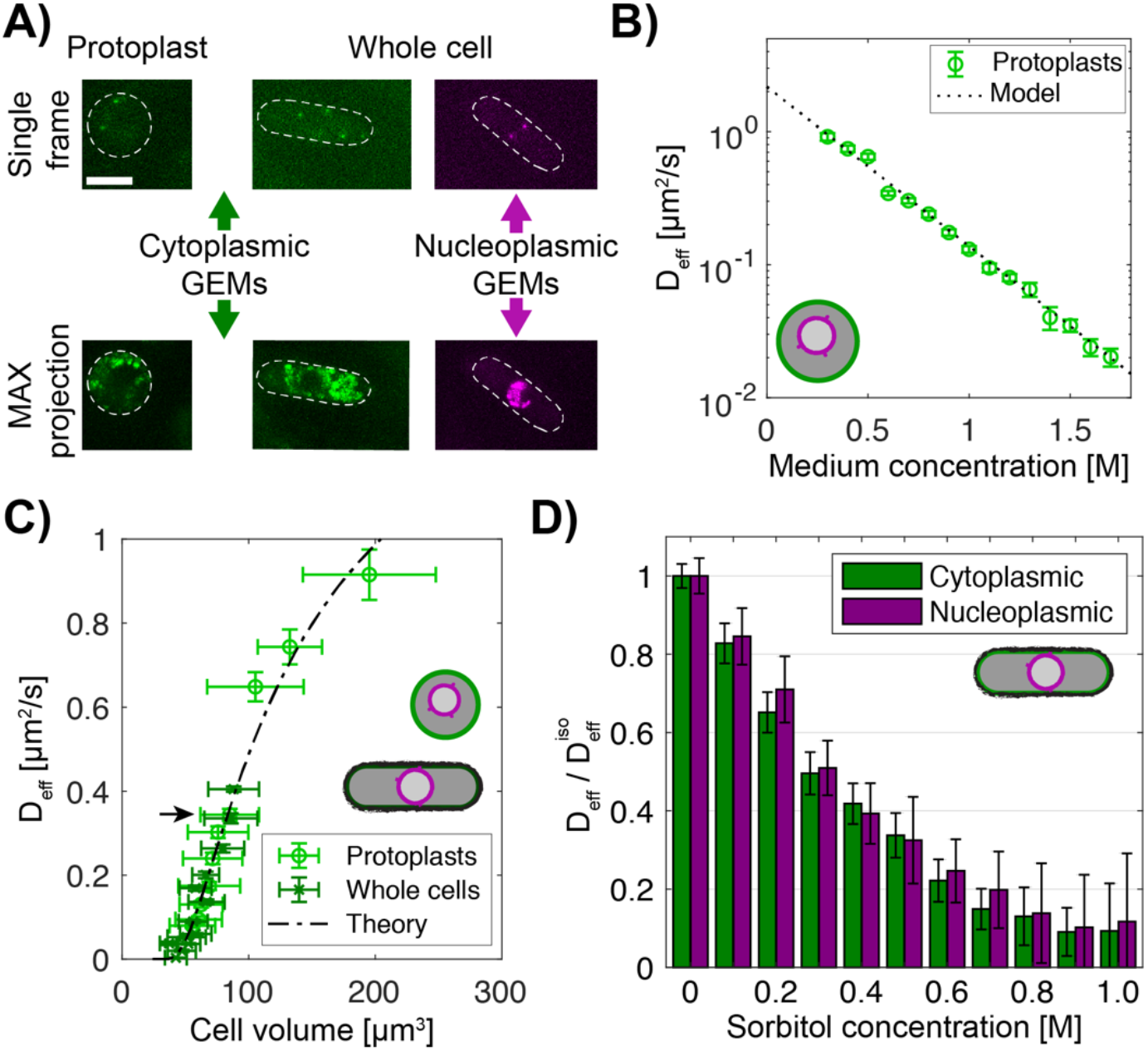
Macromolecular crowding is affected similarly in the nucleus and cytoplasm under osmotic shocks. **(A)** Images of protoplasts (left) and whole cells (right) expressing cytoplasmic 40 nm GEMs and nucleoplasmic 40 nm GEMs. Top, single time point image; bottom, maximum projection of 100 frames. Dashed lines, the cell boundary. Scale bar = 5 μm. **(B)** Effective diffusion coefficient of cytoplasmic GEMs (mean ± SEM) in protoplasts shifted to various sorbitol concentrations in the medium. Dashed lines, predictions of Phillies’ model for diffusion with a power law *λ*=1. N_GEMs_ tracked = 4058, from at least 2 biological replicates per condition. **(C)** Effective diffusion coefficient of cytoplasmic GEMs (mean ± SEM) plotted against cell volume under hypotonic and hypertonic shock. Volumes represent mean distribution of an asynchronous culture ±STD. Dashed line, fit of Phillies’ model for self-diffusing trackers in a polymer solution. Black arrow indicates Deff for a population of cells in YE and protoplasts in isotonic condition. Protoplasts: N_GEMs_ = 3355, N_Volume_ = 2216 cells, whole cells: N_GEMs_ = 9849, N_Volume_ = 981 cells. **(D)** Effects of hyperosmotic shifts on the relative effective diffusion coefficients (mean ± SEM) of cytoplasmic and nuclear GEMs; no statistically significant difference was detected (F-test, p-value=0.90). Cytoplasm N_GEMs_ = 9365, nuclear N_GEMs_ = 3732, from at least 2 biological replicates per condition. See also Figure S4.

We compared the behaviors of the GEMs in the cytoplasm and nucleoplasm. Mean square displacement (MSD) curves showed that the cytoplasmic GEMs displayed subdiffusive motion with an anomalous diffusive exponent α~0.9 comparable to measurements in HEK293 or hPNE cells, and *S. cerevisiae* (Delarue et al., 2018; Szoradi et al., 2021) (Figure S4B). Nucleoplasmic GEMs exhibited a stronger subdiffusive behavior with α~0.8, suggesting a stronger caging effect compared to the cytoplasm. Notably, nuclear GEMs consistently exhibited significantly higher D_eff_ than cytoplasmic GEMs (D^N^_eff_ ~ 0.55 μm^2^/s and D^Cy^_eff_ ~ 0.40 μm^2^/s, Figure S4C), suggesting that the nucleoplasm is less crowded at this size scale than the cytoplasm in fission yeast. Protoplasts in isotonic conditions exhibited similar D_eff_ and *α* in the cytoplasm, showing that cytoplasmic properties probed by GEMs were not affected by removal of the cell wall (Figure 4C, black arrow; S4D and E).

Next, we determined how D_eff_ of the GEMs relates to macromolecular concentration. Because of the properties of the protoplasts as ideal osmometers, we were able to quantitatively tune macromolecule concentration in the cytoplasm by using osmotic shifts. We found that D_eff_ of the cytoplasmic GEMs in protoplasts exhibited an exponential relationship with medium concentration and hence macromolecular concentration (Figure 4B). This relationship could be fit with a Phillies’ model (Masaro and Zhu, 1999; Phillies, 1988) which uses a unique stretched exponential equation to describe a tracer particle’s self-diffusive behavior in a wide range of polymer concentrations (Methods, Figures 4B, S4F). The alignment of data from walled cells and protoplasts (Figure 4C) showed that this relationship also applied to cytGEMs analyses in walled cells. Hence, these relationships showed that D_eff_ of the GEMs can be used to estimate the concentration of macromolecules over a large range of cytoplasmic concentrations.

We then used the GEMs to determine how the nucleoplasm compares with the cytoplasm in their responses to osmotic shifts. The proportionate changes of nuclear and cellular volumes (Figure 3B-D) predicted that osmotic shifts affect the cytoplasm and nucleoplasm in similar ways. Indeed, D_eff_ and *α* of cytoplasmic and nucleoplasmic GEMs responded similarly in cells treated with varying doses of sorbitol (Figures 4D, S4B, D, E). Together, these findings showed that GEMs inform on predictable changes in the concentration of macromolecules and resultant colloid osmotic pressures within both compartments; both environments showed consistent behavior without evidence for sharp transitions in biophysical properties such as phase transitions. Therefore, these findings demonstrate that the movement of GEMs provides a highly precise and quantitative approach to measure and compare macromolecular crowding within the cytoplasm and nucleoplasm.

### Inhibition of nuclear export causes an increase in the N/C ratio

An important prediction is that changes in the balance of colloid osmotic pressures between the nucleoplasm and cytoplasm would lead to a predictable change in the N/C ratio. It has been previously reported that inhibition of nuclear export leads to an increase in nuclear size, either through treatment with a drug leptomycin B (LMB, an inhibitor of the Crm1 exportin) or through mutants affecting the nuclear transport machinery (Kudo et al., 1999; Kume et al., 2017; Neumann and Nurse, 2007; Yoshida and Horinouchi, 1999). However, this effect has not been quantitatively characterized over time. We tested whether this N/C ratio increase can be explained by the accumulation of osmotically active macromolecules in the nucleus when nuclear export is impaired. LMB has been shown for instance to cause the redistribution of a subset of proteins in Xenopus oocytes (Wühr et al., 2015). In contrast to the previous studies that examined effects of LMB only after many hours of treatment (Kudo et al., 1998; Kume et al., 2019; Neumann and Nurse, 2007; Nishi et al., 1994), we examined the acute effects of LMB treatment in a time course, tracking both individual cells (Figures 5A-E) and asynchronous populations (Figures S5A-D). Upon LMB treatment, interphase cells continued to grow at a similar rate as untreated cells, but their nucleus grew relatively faster (Figures 5A-C). This growth pattern led to a progressive increase in the N/C ratio, with a 6% increase at 15 min and a 16% increase by 60 min, causing the average N/C ratio to rise from 8% to 9% (Figures 5D-E).

**Figure 5.**
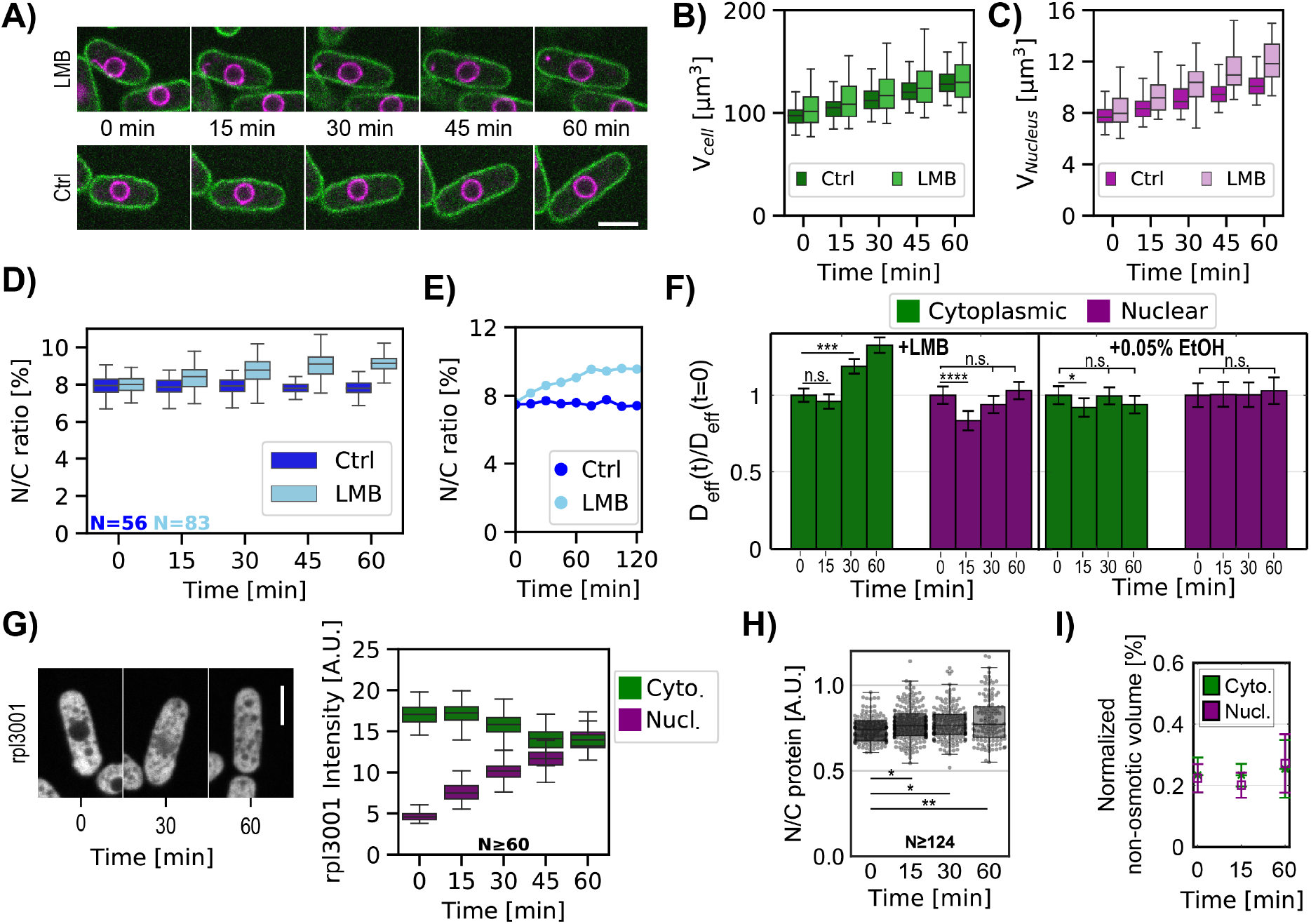
Inhibition of nuclear export rapidly leads to an increase in the N/C ratio and changes in crowding. **(A)** Individual cells expressing plasma membrane and nuclear markers were imaged in time upon treatment with LMB or control (Ctrl). Images show a mid-focal plane of plasma membrane (green) and nuclear membrane (purple) treated with LMB (top) or not (Ctrl, bottom) over time (min). **(B)** Cell volumes were measured from 3D images. **(C)** Same as (B) for nuclear volume. **(D)** Box and whisker plot of N/C ratio of individual cells treated with LMB (cyan) and control condition (blue) at t=0 min and followed by time lapse microscopy. **(E)** N/C ratio dynamics of representative individual cells extracted from (D). **(F)** Cells expressing cytoplasmic or nuclear GEMs were treated with LMB or control (0.05% ethanol) and were imaged for GEMs diffusion over time. Bar graphs show the relative changes in mean effective diffusion coefficients ± SEM for cytoplasmic (green) and nucleoplasmic (purple) GEMs in cells treated with LMB (left panel) and ethanol control (right panel). Statistical differences compared with Mann-Whitney U test. **(A-F)** From at least 3 biological replicates. **(G)** Cells expressing chromosomally tagged proteins that mark the large ribosomal subunit (Rpl 3001-GFP) were treated with LMB and imaged over time. Mid focal plane confocal images and quantitation of their relative fluorescence intensities are displayed. From 3 biological replicates. **(H)** Cells were stained for total protein using RNAse and FITC dye, plots indicate ratios of FITC intensities in nuclear and cytoplasmic regions. Statistical differences compared with Kruskal-Wallis test, from at 3 biological replicates. **(I)** Normalized non-osmotic volume over time for cells and their nuclei in protoplasts treated with LMB. Scale bar = 5 μm. See also Figure S5 and S6.

We used multiple assays to quantitatively test whether the increase in the N/C ratio was due to accumulation of macromolecules in the nucleus. First, we used the GEMs-based nanorheology to assess nuclear macromolecular crowding. D_eff_ of nuclear GEMs significantly decreased at 15 min of LMB treatment, indicating a rise in crowding in the nucleoplasm (Figure 5F). However, the D_eff_ of the nuclear GEMs subsequently increased at 30 min but returned to a normal level at 60 min, possibly due to adaptation behavior such as increasing nuclear size. Interestingly, measurements of cytoplasmic GEMs revealed a different pattern: the cytGEMs D_eff_ did not change significantly at 15 min, but increased progressively at 30 and 60 min. This result suggested that the cytoplasm became progressively less crowded (equivalent to 4% and 11% dilution at 30 and 60 min, respectively). Thus, these findings indicated both increased colloid osmotic pressure in the nucleus and decreased pressure in the cytoplasm contribute to nuclear swelling.

Second, we characterized the subcellular localization of ribosomal subunits. Ribosomes and their subunits are major components of biomass and contributors to macromolecular crowding in the cytoplasm (Delarue et al., 2018; Warner, 1999). LMB inhibits the export of large ribosomal subunit proteins, which are transported into the nucleus to be assembled into pre-60S particles and are then normally exported from the nucleus in a Crm1-dependent manner (Aitchison and Rout, 2000; Ho et al., 2000). Cells expressing large ribosomal subunits Rpl3001 and Rpl2401 tagged with the GFP (Knapp et al., 2019) displayed little detectable fluorescence intensity in the nucleus without LMB (Figures 5G, S5E). Upon LMB treatment, fluorescence intensities of both increased in the nucleus at 15 min and were nearly equivalent to cytoplasmic values at 60 min. In contrast, a small ribosomal subunit protein Rps2-GFP (Knapp et al., 2019), which was not expected to be affected by LMB (Aitchison and Rout, 2000), showed little accumulation in the nucleus under LMB treatment (Figure S5F). The cytoplasmic intensity of all three ribosomal markers decreased slightly by 45 min (Figures 5G, S5F; right panel), which may be due to ribosomal turnover and/or impaired biogenesis. Thus, these large ribosomal subunits illustrate how LMB causes redistribution of certain proteins that likely contribute to increased colloid osmotic pressure in the nucleus.

Third, to quantify the levels of total proteins, we fixed and stained cells treated with RNAse with the fluorescent dye fluorescein isothiocyanate (FITC) and analyzed their fluorescence intensities (Knapp et al., 2019; Kume et al., 2017; Odermatt et al., 2021). FITC staining in control cells suggested that the nucleus had lower protein concentration than in the cytoplasm (Figure S5F) (Odermatt et al., 2021). Using this approach, we showed that LMB-treated cells exhibited an ~8% increase in the ratio of nuclear to cytoplasmic protein concentration compared to control cells, indicating a relative accumulation of protein in the nucleus (Figures 5H, S5F). To test whether this magnitude of accumulation is consistent with the observed increase in nuclear size, we simulated the effects of redistributing solutes into the nucleus (Appendix Figure B1). Modeling predicted that the import of 10^−18^ moles of solute into the nucleus would generate a 1% increase in the N/C ratio. An 8% redistribution in solute content from the cytoplasm to the nucleoplasm yielded a similar N/C ratio increase as seen experimentally. Thus, the observed ~ 8% increase of protein in the nucleus is of the correct magnitude to explain quantitatively the increase in nuclear size.

Fourth, we determined whether LMB affected the distribution of normalized non-osmotic volumes *v_b_*. The size of the nucleus depends not only upon colloid osmotic pressure, but also on normalized non-osmotic volumes in the nucleus and cytoplasm 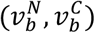. BVH plots (Figures 2D, E) indicated that in control cells the normalized non-osmotic volumes in the cytoplasm and nucleoplasm were both ~25% 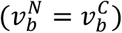. To measure the effects of LMB treatment on 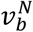 and 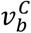, we subjected protoplasts treated with LMB to osmotic shifts. As expected, LMB caused an increase in the N/C ratio in protoplasts like that seen in whole cells (Figure S6A-D). As with control cells, osmotic shocks did not further alter the N/C ratio (Figure S6E); for instance, after 60 min of LMB treatment, protoplasts maintained an elevated N/C ratio of 10% over a range of hypoosmotic and hyperosmotic conditions (Figure S6F). Analyses of these data showed that although the LMB treatment caused an increase in the absolute amount of dry mass in the nucleus, the nucleus also expanded in volume, so that the values for non-osmotic volumes that are normalized for size remained constant at ~25% (Figure 5I). These results suggest that the accumulation of dry mass in the nucleus was compensated by expansion of nuclear volume, so that *v_b_* is maintained in both compartments.

In summary, we established that inhibiting nuclear export with LMB produced a progressive increase in the N/C ratio. This rise was accompanied by a transient increase in crowding in the nucleus and a progressive decrease in crowding in the cytoplasm, nuclear accumulation of a subset of cellular components, and compensatory changes in nuclear size.

### Protein synthesis inhibition does not alter the N/C ratio

Another way to globally perturb macromolecular levels is by inhibiting protein synthesis. As LMB treatment disrupted ribosomal biogenesis (Figure 5G, S5E), we tested whether inhibition of translation itself would alter the N/C ratio. We analyzed cells treated with 50 mg/mL cycloheximide (Polanshek, 1977) (Figure 6A). At this relatively low dosage, interphase cells continued to grow in volume but at slower rates (Figures 6B). The nuclei also grew at the same slower rate, maintaining the N/C ratio (Kume et al., 2017) (Figures 6C, D, E). This maintenance of the N/C ratio over time was also observed in asynchronous cell populations (Figures S7A-D). GEMs analyses revealed that D_eff_ of nuclear and cytoplasmic GEMs increased proportionally (Figure 6F). FITC staining indicated a progressive decrease in concentration of total protein in both the nucleus and cytoplasm (Figures 6G, H), leading to a ~30% decrease in total protein concentration after 1 h of CHX treatment.

**Figure 6.**
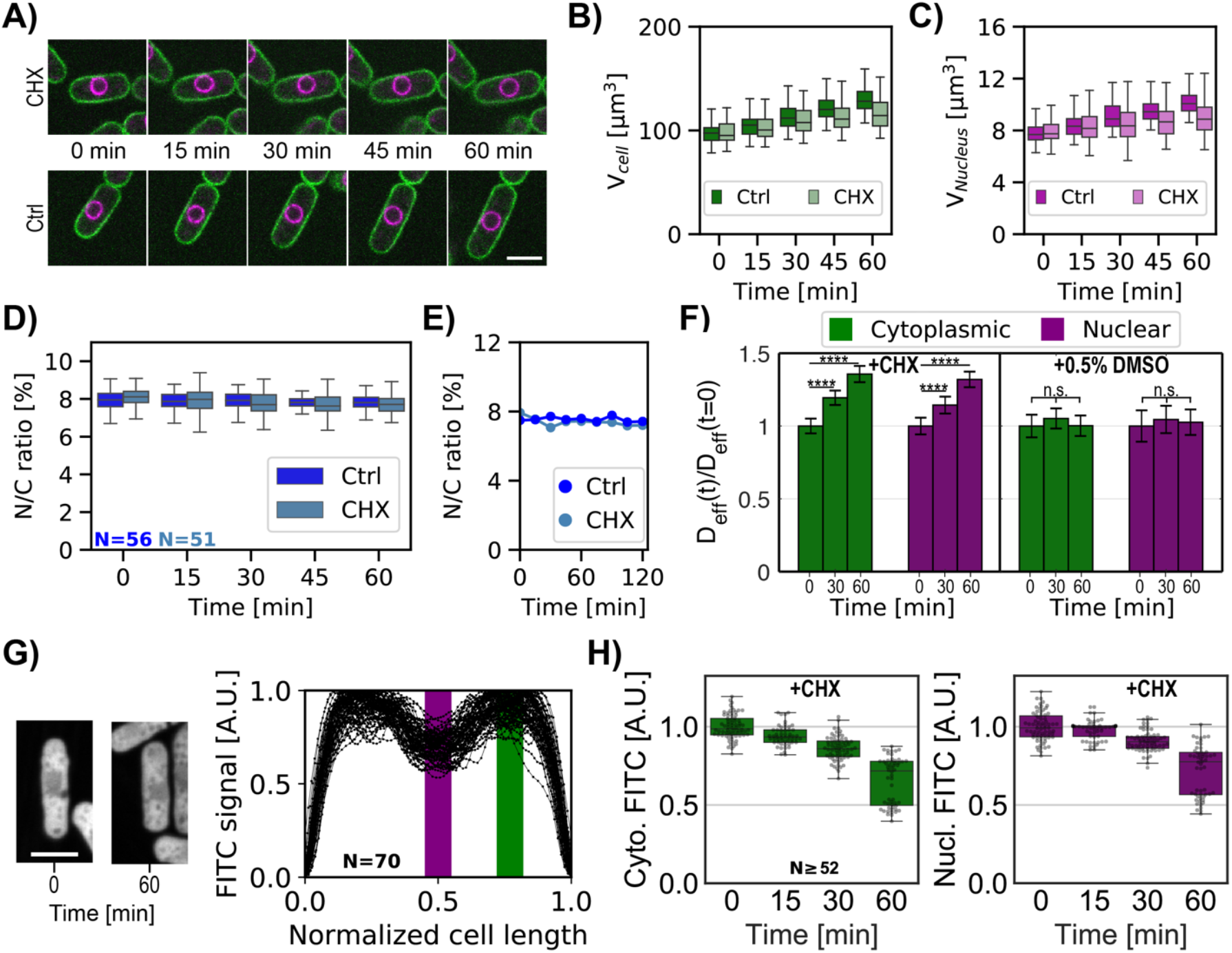
Inhibition of protein synthesis is accompanied by a similar decrease in nucleo-cytoplasmic crowding and does not perturb the N/C ratio. **(A)** Overlay of the plasma membrane (green) and nuclear membrane (purple) of whole cells middle plane over time treated with cycloheximide (CHX, top) or not (Ctrl, bottom). **(B)** Single whole-cell volume dynamics treated with CHX or not (Ctrl). **(C)** Same as (B) for single-nucleus volume dynamics treated with CHX or not (Ctrl). **(D)** Individual whole cells N/C ratio dynamics treated with CHX or not (Ctrl). **(E)** Single whole-cell N/C ratio dynamics extracted from (D) for each condition. **(F)** Relative cytoplasmic (green) and nucleoplasmic (purple) GEM effective diffusion dynamics for cells treated with CHX (left panel) or only the drug buffer for control (0.5% dimethyl sulfoxide (DMSO), right panel). Statistical differences compared with Mann-Whitney U test. **(G)** Left, confocal middle plane image of FITC-stained cells treated with RNAse to remove RNA staining, before (0 min) or after CHX treatment for 60 min. Right, quantification of FITC fluorescence along the long cell axis, normalized by cell length. Purple and green boxes indicate the positions of nuclear and cytoplasmic signals used to build the graph in (H). **(H)** Cytoplasmic (green, left) and nucleoplasmic (purple, right) protein signals for the same cells over time under CHX treatment decrease similarly. From 3 biological replicates Scale bar = 5 μm. See also Figure S7. **(A-H)** From at least 3 biological replicates.

These findings demonstrate that the proportionate dilution of macromolecular components in both compartments leads to a proportionate decrease in growth rates and maintenance of the N/C ratio. These experimental results strengthen our model of a nucleus behaving like an ideal osmometer for which a similar decrease in osmotically active particles in both sides of the nuclear envelope leads to a constant N/C ratio.

### N/C ratio homeostasis can be explained by an osmotic model for cell and nuclear growth

The N/C ratio is maintained with little variability, with a coefficient of variation of the N/C ratio of ~ 0.1 (Figure 7A, WT). The ratio is robustly maintained throughout the course of cell growth during the cell cycle (Figure 5D) (Jorgensen et al., 2007; Neumann and Nurse, 2007), indicating that nuclear volume normally grows at the same rate as the volume of the cytoplasm. Like many other cell types, the growth rate of fission yeast cells is largely exponential in character, such that large cells grow faster than smaller ones (Knapp et al., 2019; Pickering et al., 2019; Tzur et al., 2009). One basis for this size dependence is thought to be due to the scaling of active ribosome number in the cytoplasm to cell size. The low variability of the N/C ratio suggests that it may be maintained by a homeostasis mechanism, so that cells with aberrant N/C ratio correct their nuclear size. Indeed, it was recently reported that *S. pombe* cells exhibit homeostasis behavior to maintain nuclear scaling (Cantwell and Nurse, 2019a; Neumann and Nurse, 2007). We sought to quantify this homeostasis behavior and to test whether the N/C ratio correction could be explained by a passive osmotic model or whether an additional active feedback mechanism needs to be invoked.

**Figure 7.**
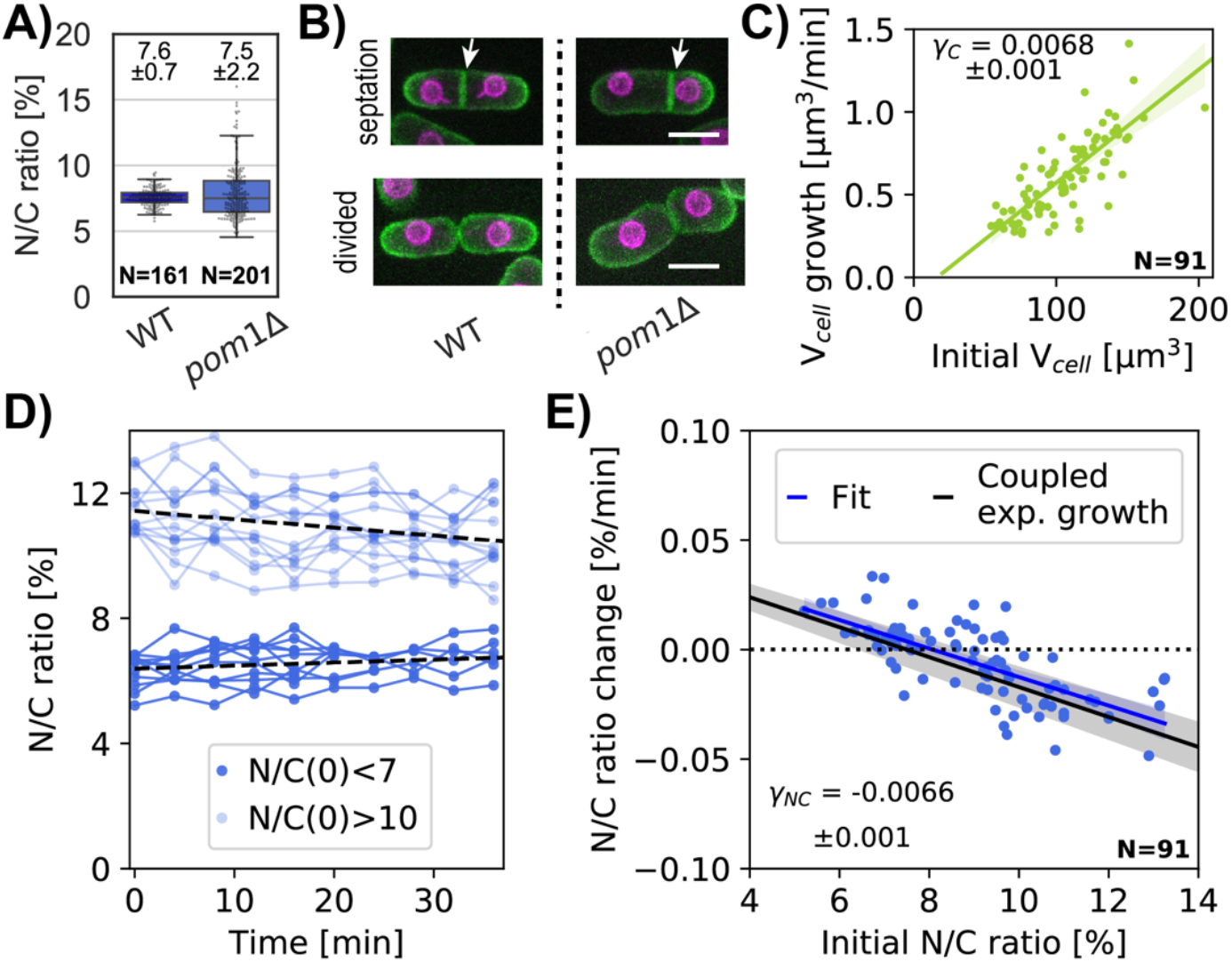
Correction of aberrant N/C ratio follows a passive nuclear growth model correlated with cell size. **(A)** Asynchronous WT and *pom1*Δ whole cells N/C ratio (mean ±STD) in growth medium, from 1 biological replicate. **(B)** Z-sum projection overlay of the plasma membrane (green) and nuclear membrane (purple) of representative cells at septation (top) and divided (bottom) for WT (left) and *pom1*Δ(right). White arrow, septum location in the middle of WT cells and decentered for *pom1*Δ cells leading to asymmetric cell division. Scale bar = 5 μm. **(C)** Cellular growth rate as a function of a cell’s initial volume. Linear regression is shown in green with a slope *γ*_C_. **(D)** N/C ratio over time for selected cells with low (light blue) or high (dark blue) initial N/C ratio. Dashed lines, linear regression for each cohort of cells. **(E)** N/C ratio change over time as a function of the initial N/C ratio. Experimental data (blue dots), linear fit (blue line), and predicted passive homeostasis N/C ratio behavior (black line) assuming N/C ratio =7.5% at equilibrium from (B) and cell growth rate *γ*_C_ from (C). See also Figure S8. **(C-E)** From 2 biological replicates.

We measured homeostasis behavior in *pom1*Δ mutant cells, which display variable N/C ratios because of asymmetric cell division (Bahler and Pringle, 1998; Cantwell and Nurse, 2019a). Time lapse images showed that these cells exhibited normal mitosis that produced two equally sized nuclei, but often placed their division septum asymmetrically, yielding daughters that were born with either too low or too high N/C ratios (Figures 7A-B, S8A-B). Consistent with this, an asynchronous population of *pom1*Δ cells displayed the same average N/C ratio as wildtype but with a ~3-fold larger standard deviation (Figure 7A).

We tracked cell and nuclear growth in these cells with abnormal N/C ratio as they grew during interphase (Methods; Figures S8C-E). Under these conditions, cells exhibited an exponential growth rate of *γ_C_* ≈ 0.006 μm^3^/min, corresponding to an expected doubling time of ~115 min (Figure 7C). Cells born with abnormally large N/C ratios (>10) or abnormally small ratios (<7) gradually corrected their N/C ratio over time (Figures 7D, S8F), consistent with previous findings (Cantwell and Nurse, 2019a). Nuclei in cells with high N/C ratios grew slower than those in cells with normal N/C ratios, while nuclei in cells with low N/C ratios grew faster. In contrast, cells with near normal N/C ratios showed little change in this ratio over time. We detected an inverse relationship between the initial ratio and the rate of correction (Figure 7E, blue dots). By fitting the N/C ratio change for a population of *pom1*Δ cells, we quantified the N/C ratio growth rate as a function of the initial N/C ratio (Figure 7E, blue line).

A previous paper proposed a model for N/C ratio homeostasis based upon an active feedback mechanism (Cantwell and Nurse, 2019a). We tested whether this N/C ratio correction could be explained by a passive osmotic model without feedback. We built upon our simple osmotic model to incorporate dynamic growth (See Methods and Appendix B.2 for detailed derivations). We assumed that volume growth is driven by the rate of biosynthesis of cellular components (Altenburg et al., 2019; Knapp et al., 2019; Midtvedt et al., 2019; Odermatt et al., 2021), likely dependent, in part, on the number of active ribosomes in the cytoplasm. The growth of the cytoplasm is driven by the biosynthesis of osmotically-active macromolecules targeted for the cytoplasm. Similarly, the growth of the nucleus is driven by the biosynthesis of macromolecules in the cytoplasm that are transported into the nucleus; assuming that nuclear transport is not limiting, the rate of nuclear growth may thus also scale with the volume of the cytoplasm. The balance of colloid osmotic forces in each compartment determines the cell and nuclear volumes. We also assumed that the rate of synthesis of nuclear components is a fixed percentage of total synthesis rate (e.g., 7.5%). Thus, assuming that the nucleus is an ideal osmometer, the percentage of total synthesis rate that goes into the nucleus versus cytoplasm is what ultimately sets the N/C ratio at equilibrium.

By assuming exponential cell growth, we can compute the change in N/C ratio over time such that:

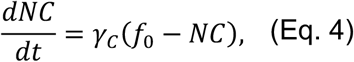

where *f*_0_ is a constant and represents the fraction of osmotically active particles transported into the nucleus and *γ_C_* is the exponential cellular growth rate. Our measurements gave us access to every parameter in Eq. 4 with no free parameters.

The model predicts that in cells with an altered N/C ratio, the N/C ratio returns to *f*_0_ over time, hence *f*_0_ = 7.5 ± 0.7 % (Figure 7A, Appendix Figure B2). We used this relationship to determine the homeostasis behavior of the N/C ratio (Figure 7E, black line). Our experimental data were an excellent fit for this prediction using the measured parameters (*γ_C_* and N/C ratio at equilibrium) (Figure 7E; blue versus black lines). This result suggests that maintaining the synthesis rate of nuclear contents at 7.5% of the cellular synthesis rate is sufficient to explain not only the coupling of nuclear growth rates with cell growth in normal growth, but also accounts for homeostasis correction when the N/C ratio is altered. The correction rates are on the time scales of growth rates; therefore, the model predicts that full correction of significant N/C alterations requires periods of multiple generation times, and that under certain circumstances this correction is more rapid assuming exponential growth dynamics rather than linear growth (Appendix Figure B2).

## Discussion

Here, we provide a quantitative model for nuclear size control based upon osmotic forces. This model, which has zero free parameters, postulates that nuclear size is dictated in part by the number of osmotically active molecules in the nucleus and cytoplasm that cannot readily diffuse through the nuclear membranes (Figure 1; see Appendix). These molecules, which include large proteins, RNAs, metabolites and other large molecules (>30 kDa), produce colloid osmotic pressure on the nuclear envelope to expand nuclear volume to a predicted size at steady state. Another potential parameter is membrane tension of the nuclear envelope, as determined by its ability to expand under pressure (Figure 1). In fission yeast, we determined that the nucleus readily changes in volume under osmotic perturbations, thus behaving as a near-ideal osmometer (Figure 2E); this behavior indicates that contributions of membrane tension of the nuclear envelope on nuclear size are negligible (or very small). The N/C ratio is then set as the ratio of nuclear to cytoplasmic solutes (Eq. 3); in the case of fission yeast, the number of these nuclear solutes is predicted to be about 8% of the total in the cell, giving rise to an average N/C ratio of 8%. Therefore, using this system, we confirm and define quantitatively the primary contributions of osmotic pressures to nuclear size control.

This physical model explains why the N/C ratio is so robustly maintained in the vast majority of mutant and physiological conditions (Cantwell and Nurse, 2019c). The N/C ratio arises because the cell globally maintains the relative quantities of nuclear to cytoplasmic solutes by protein expression and transport. During cell growth, the nucleus grows at the same rate as the cell because its growth is driven by the synthesis and transport of the nuclear macromolecules that contribute to osmotic pressure. This growth mechanism of the nucleus also explains the homeostasis behavior observed when the N/C ratio is too high or low (Figure 7). (Cantwell and Nurse, 2019a). The gradual correction of the N/C ratio by growth is reminiscent to the “adder behavior” for cell size homeostasis (Campos et al., 2014; Taheri-Araghi et al., 2015).

This proposed view suggests that the primary function of nuclear size control is perhaps not to specify a certain size, but to maintain healthy levels of macromolecular crowding in the nucleoplasm (Ellis, 2001). Our osmotic perturbations and GEMs measurements showed that the nuclear and cytoplasmic environments, which contain quite different components, nevertheless have similar degrees of mesoscale macromolecular crowding and normalized non-osmotic volumes *v_b_*. Even after long exposure to LMB, when cells exhibit a significative higher N/C ratio (Figure 5D), the normalized non-osmotic volumes of the nucleus and crowding in the nucleoplasm is similar to those in control cells (Figure 5F, 5I). The osmotic nature of nuclear size control thus allows nucleoplasm and cytoplasm to stay in balance through not only synthesis and transport but also by osmotic control of nuclear volume.

Our study provides a critical quantitative confirmation of proposed colloid osmotic pressure effects in cells, which is caused by the crowding of macromolecules (Harding and Feldherr, 1958, 1959; Mitchison, 2019). As the *in vivo* relevance of such pressures has not yet been firmly established, our results using GEMs, osmotic shifts and fluorescent intensity measurements constitute a strong experimental demonstration of the importance of these forces. Although similar osmotic-based models for nuclear size control have been proposed (Churney, 1942; Kim et al., 2016), our studies benefited from the accurate measurements and quantitative analysis possible in this model cell type.

This osmotic-based mechanism is likely the primary factor in nuclear scaling mechanisms in mammalian and other cell types. Osmotic shift experiments in mammalian chondrocytes cells (Finan et al., 2009) reveal that the nucleus in these cells is not an ideal osmometer, but is restricted from swelling due to hypoosmotic shocks because of nuclear membrane tension. Indeed, the BVH plot in this paper (Figure 1A of (Finan et al., 2009)) can be fitted with our model assuming N^N^ = 4.10^−16^ mol and a substantial nuclear membrane tension of σ^N^ = 0.02 N/m. This additional nuclear membrane tension may be due to nuclear lamina, peri-nuclear actin and/or perinuclear chromatin forces acting on the nuclear envelope, which may be absent or reduced in fission yeast (Edens et al., 2017; Newport et al., 1990; Schreiner et al., 2015). Thus, it is likely that osmotic forces act in concert with other mechanical elements to set nuclear size in these more complex systems. We note that models similar to ours can even account for bacterial nucleoid size scaling (Gray et al., 2019), where instead of a nuclear envelope tension term, a partition coefficient establishes the equilibrium concentration of molecules between the cytoplasm and the nucleoid/nucleus. Elucidation of mechanisms rooted in physics promise to give new insights into the range of nuclear shapes and sizes seen during development as well in diseases and aging (Foraker, 1954; Karoutas and Akhtar, 2021; Roubinet et al., 2021; Zink et al., 2004). Approaches such as osmotic shifts and nanorheology will allow for future investigation of similar osmotic mechanisms responsible for size control of the nucleus and other organelles.

## Materials and Methods

### Yeast strains and media

The *Schizosaccharomyces pombe* strains used in this study are listed in Supplemental Table S1. In general, fission yeast cells were grown in rich medium YES 225 (#2011, Sunrise Science Production) at 30 °C. Strains carrying GEM expression vectors were grown in EMM3S – Edinburgh Minimum Media (#4110-32, MP Biomedicals) supplemented with 0.225 g/L of uracil, histidine, and adenine as well as 0.05 μg/mL of thiamine (#U0750, #H8000, #A9126, #T4625, Sigma-Aldrich).

### Microscopy

Cells were imaged on a Ti-Eclipse inverted microscope (Nikon Instruments) with a spinning-disk confocal system (Yokogawa CSU-10) that includes 488 nm and 541 nm laser illumination and emission filters 525±25 nm and 600±25 nm respectively, a 60X (NA: 1.4) objective, and an EM-CCD camera (Hamamatsu, C9100-13). These components were controlled with μManager v. 1.41 (Edelstein et al., 2010, 2014). Temperature was maintained by a black panel cage incubation system (#748-3040, OkoLab).

For imaging of GEMs, live cells were imaged with a TIRF Diskovery system (Andor) with a Ti-Eclipse inverted microscope stand (Nikon Instruments), 488 nm laser illumination, a 60X TIRF oil objective (NA:1.49, oil DIC N2) (#MRD01691, Nikon), and an EM-CCD camera (Ixon Ultra 888, Andor), controlled with μManager v. 1.41 (Edelstein et al., 2010, 2014). Temperature was maintained by a black panel cage incubation system (#748-3040, OkoLab).

For most live cell imaging, cells were mounted in μ-Slide VI 0.4 channel slides (#80606, Ibidi – 6 channels slide, channel height 0.4 mm, length 17 mm, and width 3.8 mm, tissue culture treated and sterilized). The μ-Slide channel was coated by pre-incubation with 100 μg/mL of lectin (#L1395, Sigma) for 15 min at room temperature, and then washed with medium. Cells in liquid culture were introduced into the chamber for 3 min and then washed three times with medium to remove non-adhered cells. As certain drugs may adhere to polymer slide material in the conventional chambers, μ-Slide chambers with glass bottoms (#80607, Ibidi – 6 channels slide, channel height 0.54 mm, length 17 mm and width 3.8 mm, D263M Schott glass and sterilized) were used for the drug treatments.

### 3D volume measurements

Nuclear and cell volumes were measured in living fission yeast cells expressing a nuclear membrane marker (Ish1-GFP, (Expósito-Serrano et al., 2020)) and a plasma membrane marker (mCherry-Psy1,(Kashiwazaki et al., 2011)) using a semi-automated 3D segmentation approach. Z stack images (0.5 μm z-slices) that covered the entire cell (for a total of ~20 slices) were obtained using spinning disk confocal microscopy. The 3D volumes were segmented using an ImageJ 3D image segmentation tool LimeSeg (Machado et al., 2019; Schneider et al., 2012) with these parameters:

For cells: run(“Sphere Seg”, “d_0=3.0 f_pressure=0.016 z_scale=4.5 range_in_d0_units=3.0 color=51,153,0 samecell=false show3d=false numberofintegrationstep=-1 realxypixelsize=0. 111”); For nuclei: run(“Sphere Seg”, “d_0=2.0 f_pressure=0.016 z_scale=4.5 range_in_d0_units=2.0 color=51,153,0 samecell=false show3d=false numberofintegrationstep=-1 realxypixelsize=0. 111”);

After each 3D analysis converged, segmentation results were confirmed using a 2D result. If there was a discrepancy, additional analyses on individual cells were used, with multiple circular regions of interest if necessary. Volume data were analyzed with a custom Python script on Jupiter Notebook 5.5.0. In general, experiments are representative of at least 2 biological replicates with independent data sets as described in the figure legends.

### Protoplast preparation

*S. pombe* cells were inoculated from fresh agar plates into YES 225 or EMM3S liquid cultures and grown at 30 °C for about 20 h into exponential phase (OD_600_ = 0.2-0.3). Ten milliliters of cells were harvested by centrifugation 2 min at 0.4 rcf, washed two times with SCS buffer (20 mM citrate buffer, 1 M D-sorbitol, pH=5.8), resuspended in 1 mL of SCS buffer with 0.1 g/mL Lallzyme (#EL011-2240-15, Lallemand), and incubated with gentle shaking for 10 min at 37 °C in the dark (Flor-Parra et al., 2014b). The resulting protoplasts were gently washed three times in YES 225 or EMM3S with 0.4 M D-sorbitol, using centrifugation for 2 min at 0.4 rcf between washes. After the last wash, 900 μL of supernatant were removed, and the protoplasts in the pellet were gently resuspended in the remaining ~100 μL of solution. The resultant protoplasts were introduced into a lectin-coated μ-Slide VI 0.4 channel slide for imaging.

### Osmotic shocks

Fission cells or protoplasts were loaded in a lectin-treated μ-Slide VI 0.4 channel slide and maintained at 30 °C. After 5 min of incubation, cells were washed three times with their respective initial buffer (isotonic condition). Cells were imaged first in their initial buffer (isotonic condition). Then, hyper or hypotonic medium was introduced into the channel with three washes. For hypotonic medium YES 225 was diluted with sterile water. The same individual cells were then imaged (within 1 min of the osmotic shift) using the same parameters. To minimize the effect of volume adaptation response to osmotic shock, we assayed cells within 1 min of the osmotic shift and performed most of our experiments using cells in *gpd1*Δ mutant background that is defective in this response (Figure S1 B and C) (Hohmann, 2002; Minc et al., 2009).

### Diffusion imaging and analysis of GEMs

For cytoplasmic 40 nm GEMs, Pfv encapsulin-mSapphire was expressed in fission yeast cells carrying the multicopy thiamine-regulated plasmid pREP41X-Pfv-mSapphire (Delarue et al., 2018; Molines et al., 2020). For nuclear 40 nm GEMs, NLS-Pfv-mSapphire was inserted pREP41X (Szoradi et al., 2021). The expression of these constructs was under the control of the thiamine repressible *nmt41* promoter (Maundrell, 1990). Cells were grown using a protocol that produced appropriate, reproducible expression levels of the GEMs: cells carrying these plasmids were grown from a frozen stock on EMM3S - LEU plates without thiamine for 2-3 days at 30 °C and stored at room temperature for 1-2 days to induce expression. Cells were then inoculated in liquid EMM3S -LEU with 0.1 μg/mL of thiamine (#T4625-25G, Sigma Aldrich) for partial repression of the *nmt41* promoter and grown for one day at 30 °C to exponential phase.

Cells in lectin-treated μ-Slide VI 0.4 channel slides (#80606, Ibidi) were imaged in fields of 250×250 pixels or smaller using highly inclined laser beam illumination at 100 Hz for 10 s. GEMs were tracked with the ImageJ Particle Tracker 2D-3D tracking algorithm from MosaicSuite (Sbalzarini and Koumoutsakos, 2005) with the following parameters:

> run(“Particle Tracker 2D/3D”, “radius=3 cutoff=0 per/abs=0.03 link=1 displacement=6 dynamics=Brownian”).

The analyses of the GEMs tracks were like those described in (Delarue et al, 2018), with methods to compute mean square displacement (MSD) using MATLAB (MATLAB_R2018, MathWorks). The effective diffusion coefficient *D_eff_* was obtained by fitting the first 10 time points of the MSD curve (*MSD_truncated_*) to the canonical 2D diffusion law for Brownian motion: *MSD_truncated_*(*τ*) = 4*D_eff_τ*. In general, experiments are representative of at least 2 biological replicates with independent data sets as described in the figure legends.

### LMB treatment

A stock solution of 0.1 mM LMB (#87081-35-4, Alfa Aesar) in ethanol (#BP2818-500, Fisher BioReagents) was prepared. The final concentration of 25 ng/mL in YES 225 contained 2.3 uL of the stock solution and 5 mL of cell culture. For imaging individual cells over time, exponential phase cells were placed in a μ-Slide VI 0.5 glass bottom channel slide (#80607, Ibidi). Cells where washed three times with a solution of YES 225 + 25 ng/mL LMB and then imaged. For measurements of a population of cells over time, exponential-phase cells were incubated with the drug at 30 °C with shaking. At each time point, 1 mL of the cell culture was harvested and centrifuged for 2 min at 0.4 rcf. One microliter of the pellet was spread on an 1% agarose (#16500500, Invitrogen) pad (with no drug added), sealed with Valap, and imaged within 5 min.

### Cycloheximide treatment

Cycloheximide (#C7698, Sigma-Aldrich) stock was prepared at 5 mg/mL in dimethyl sulfoxide (#67-68-5, Fisher Scientific) and stored at −20 °C. CHX was added to a final concentration of 50 μg/mL to cell cultures and samples were imaged in μ-Slide VI 0.5 glass bottom channel slides (#80607, Ibidi).

### FITC staining

Total protein was measured in individual fission yeast cells using FITC staining, similarly as described (Knapp et al., 2019; Odermatt et al., 2021). One milliliter of cell culture was fixed with 4% formaldehyde (methanol-free, #28906, Thermo Scientific, Waltham) for 60 min, washed with phosphate buffered saline (PBS) (#14190, Thermo Scientific,), and stored at 4 °C. One hundred microliters of fixed cells was treated with 0.1 mg/mL RNAse (#EN0531, Thermo Scientific) and incubated in a shaker for 2 h at 37 °C. Next, cells were washed and re-suspended in PBS and stained with 50 ng/mL FITC (#F7250, Sigma) for 30 min, washed three times, and resuspended in PBS. Cells were mounted on a 1% agarose + PBS pad and imaged in bright field and with 488 nm laser illumination via spinning disk confocal microscopy. The FITC signal was acquired in 300 nm z-step stacks that covered the entire cell volume. For each selected cell, the FITC signal intensities were measured along the long cell axis (averaged over 10 pixels in width) and normalized by cell length. The signal was corrected for background intensity and normalized by the maximum intensity along the line profile within each cell (Figures 5G, 6H, 6J). The nuclear and cytoplasmic FITC signals were defined as the sum of the signal from respectively 0.45-0.55 (middle of the cell, for the nucleus) and 0.7-0.8 (for the cytoplasm) along the normalized cell length normalized by the mean value at 0 min.

### N/C ratio homeostasis measurements

*pom1*Δ cells were grown in exponential phase in YES 225, loaded in a μ-Slide VI 0.4 channel slide (#80606, Ibidi), and imaged every 4 min for 40 min at 30 °C. The volumes of each cell and nucleus were measured over time, and growth rates were obtained by extracting the slope of a linear regression to the data over 40 min using a custom-written Python script. Data points after which cells entered nuclear division in mitosis were not included in the analysis.

### Determination of the intracellular osmolarity in *S. pombe*

For an ideal osmometer, the volume is solely determined by the balance between the outside and the intracellular concentration of osmotically active particles. As we reported that protoplasts behave like ideal osmometers (Figure 2C, D) they can therefore be used to quantify the number of osmolytes (*N^C^*) in *S. pombe*. For an ideal osmometer, *N^C^* is directly related to the cell volume (*V^C^*):

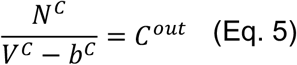

We explored the response in protoplasts volumes *V^C^* to changes in medium concentration (C^out^) for various osmotic shocks. Protoplasts were prepared in an isotonic solution and shifted in hypo or hyper conditions by the addition or removal of sorbitol in the buffer. The variation of total concentrations ΔC^out^=C^final^-C^initial^ were known. Meanwhile, variations of cells’ volume were measured before and after shocks. Since the cell non-osmotic volume b^C^ does not vary under osmotic shocks, we extracted the only unknown parameter of the equation for each cells N^C^. We found that N^C^ is linearly related to the cell volume (Figures SC-G), which means that cells keep a constant concentration of osmolytes during cell cycle. We also confirmed by analyzing various shocks such that ΔC^out^ spanned from −0.2 to 0.6 M that N^C^ does not depend on the range of osmotic shocks explored to measure it (Figure S1 C). The intracellular osmolyte concentration in *S. pombe* remained constant at a concentration of ~30.10^7^ solutes/μm^3^.

### Measurement of effective diffusivity

Under acute osmotic perturbations, cell volume changes due to the flow of water, which also affects molecular crowding and the effective GEMs diffusion. We took advantage of these quantitative measurements to assess whether the change in GEMs movements due to osmotic shocks could be explained with a physical model of diffusion in polymer solution. Phillies’ model (Masaro and Zhu, 1999; Phillies, 1988) uses a unique stretched exponential equation to describe a tracer particle self-diffusive behavior in a wide range of polymer concentrations.

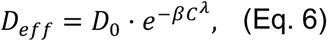

where *D*_0_ is the diffusion of the tracer particle in aqueous solution, C the concentration in polymers and *β* and *γ* are scaling parameters. D_0_ can be calculated using the Stokes-Einstein relation for a spherical particle of 40 nm diameter in water. Because protoplasts behave like ideal osmometer (Figure 2C, D) their macromolecular intracellular concentration is proportional to the medium concentration C^out^. We took advantage of this behavior to probe the variation of D_eff_ as a function of the medium concentration and found that *λ* = 1 fit our data (Figure 4B, S4F). *λ* has been found in *in vitro* experiments to depend on the molecular weight of the proteins (Banks and Fradin, 2005). Interestingly, *λ* ≈ 1 corresponds to a polymer molecular weight of 43.5 kDa close to the average protein molecular weight for Eukaryotes (~50 kDa (Milo and Phillips, 2015)) that fits our *in vivo* data. Protoplasts behave like ideal osmometers such that we can express the intracellular concentration C in the Phillies’ model as a function of the cell volume for each osmotic shock and see whether D_eff_ follows this model for which we now have only one free parameter. The model, assuming a change in intracellular concentration, is in agreement with our experimental values under acute osmotic shifts (Figure 4B). We also found that the values for D_eff_ and cell volumes measured on whole cells followed the same model (Figure 4C).

### Modeling nuclear growth and N/C ratio homeostasis

We started with a simple model for which nuclear growth was proportional to cell growth while keeping the osmotic behavior of the nucleus: nuclear volume is proportional to the number of osmotically active particles it contains. To determine the cells’ growth rate (*γ_C_*), we imaged cells at 30 °C initially at various stages of the cell cycle every 4 min for 40 min or until division happened. We plotted the change in cells volume as a function their volume at the beginning of the experiment and found a good linear correlation revealing that *S. pombe* growth is exponential with a growth rate *γ_C_* ≈ 0.006 μm^3^/min (Figure 7C) in the same range of previously reported values:

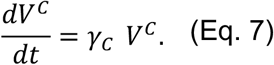

If now we assume that the nuclear growth rate is coupled (by a factor *f*_0_) to cell growth rate and maintained constant, then the change in nuclear volume can be written such that:

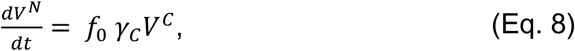

where *f*_0_ is a constant that represents the fraction of osmotically active particles synthesized by the cell that will enter the nucleus. As shown in SM Section S2.2, combining Eq. 7 and Eq. 8 results in Eq. 3 for the rate of change of the N/C ratio.

## Acknowledgments

We are grateful to members of the Chang lab, Sophie Dumont and her lab, Jane Kondev, Ariel Amir and Orna Cohen-Fix for helpful discussions. We thank KC Huang and Gant Luxton for their comments on our manuscript. Catherine Tan for building the pREP41X-Pfv-Sapphire plasmid used in this study, Rafael Daga and Yasushi Hiraoka for providing the ish1-GFP strain. T.G.F. acknowledges support under National Science Foundation grant DMS-1913093. L.J.H. acknowledges support for American Cancer Society, Pershing Square Sohn Cancer Award, Chan Zuckerberg Initiative, NIH R01 GM132447 and R37 CA240765. F.C. was supported by NIH GM05636, GM136438, NSF/BIO MCB-1638195. The authors J.L. P.RC and T.G.F. participated in the QCBNet Hackathon meeting supported by NSF under MCB-1411898.

## Author Contributions

Conceptualization: J.L., T.G.F., P.RC. and F.C.; Methodology: J.L., T.G.F. and F.C.; Software: J.L., T.G.F.; Validation: J.L., P.RC.; Formal analysis: J.L., T.G.F.; Investigations: J.L., T.G.F., P.RC.; Writing: J.L., T.G.F., and F.C.; Review editing: J.L., T.G.F., P.RC. and F.C.; Visualization: J.L., T.G.F.; Supervision: J.L., F.C.; Project administration and funding acquisition: T.G.F., F.C.

## Supplementary Material

**Table S1.**

| Yeast Strains | Source | Identifier | Figures |
| --- | --- | --- | --- |
| *h- ade6<<mCherry-psy1 ish1-GFP:kanMX ura4-D18* | This manuscript | FC3318 | 5A-E, 6A-E, 7A-B, S1A-C, S5A-D, S7, S8A-B |
| *h- ade6<<mCherry-psy1 ish1-GFP:kanMX gpd1::hphMX6 ura4-D18 ade6-* | This manuscript | FC3290 | 2A-B, 2D-E, 3, 5I, 7, S1A-C, S2C-G, S3, S6 |
| *h- gpd1::hphMX6 ade6-M216 leu1-32 ura4-D18 his3-D1* | This manuscript | FC3291 | 5H, 6G-H, S5F |
| *h? cut11-GFP:ura4+ ade6:mCherry-psy1 ura4-D18 leu1-32 ade6-M210* | This manuscript | FC3319 | S1A |
| *h- gpd1::hphMX6 pREp41X-Pfv-Sapphire leu1-32 ade6- leu1-32 ura4-D18 his7-366* | This manuscript | FC3320 | 4, S4 |
| *h- gpd1::hphMX6 pREp41X-NLS-Pfv-Sapphire leu1-32 ade6- leu1-32 ura4-D18 his7-366* | This manuscript | FC3321 | 4A, 4D, S4B-C |
| *h- pREp41X-Pfv-Sapphire ade6-M216 leu1-32 ura4-D18 his3-D1* | This manuscript | FC3289 | 5F, 6F |
| *h- pREp41X-NLS-Pfv-Sapphire leu1-32 ade6- leu1-32 ura4-D18 his7-366* | This manuscript | FC3322 | 5F, 6F |
| *h- pom1::ura4 ade6<<mCherry-psy1 ish1-GFP:kanMX* | This manuscript | FC3323 | 7A-E, S8A-F |
| *h+ rpl3001-GFP::kanR leu1-32 ura4-D18 ade6-210* | Chang Lab collection | FC3215 | 5G |
| *h- rpl2401-GFP::kanR leu1-32 ura4-D18 ade6-216* | Chang Lab collection | FC3213 | S5E |
| *h- rps2-GFP::kanR leu1-32 ura4-D18 ade6-210* | Chang Lab collection | FC3209 | S5E |
| *h+ act1p::1XE2C:HygR leu2:GFP-psy1 leu1- ura4-D18 his7-366* | This manuscript | FC3324 | S2A-B |

### Supplemental figures

**Figure S1.**
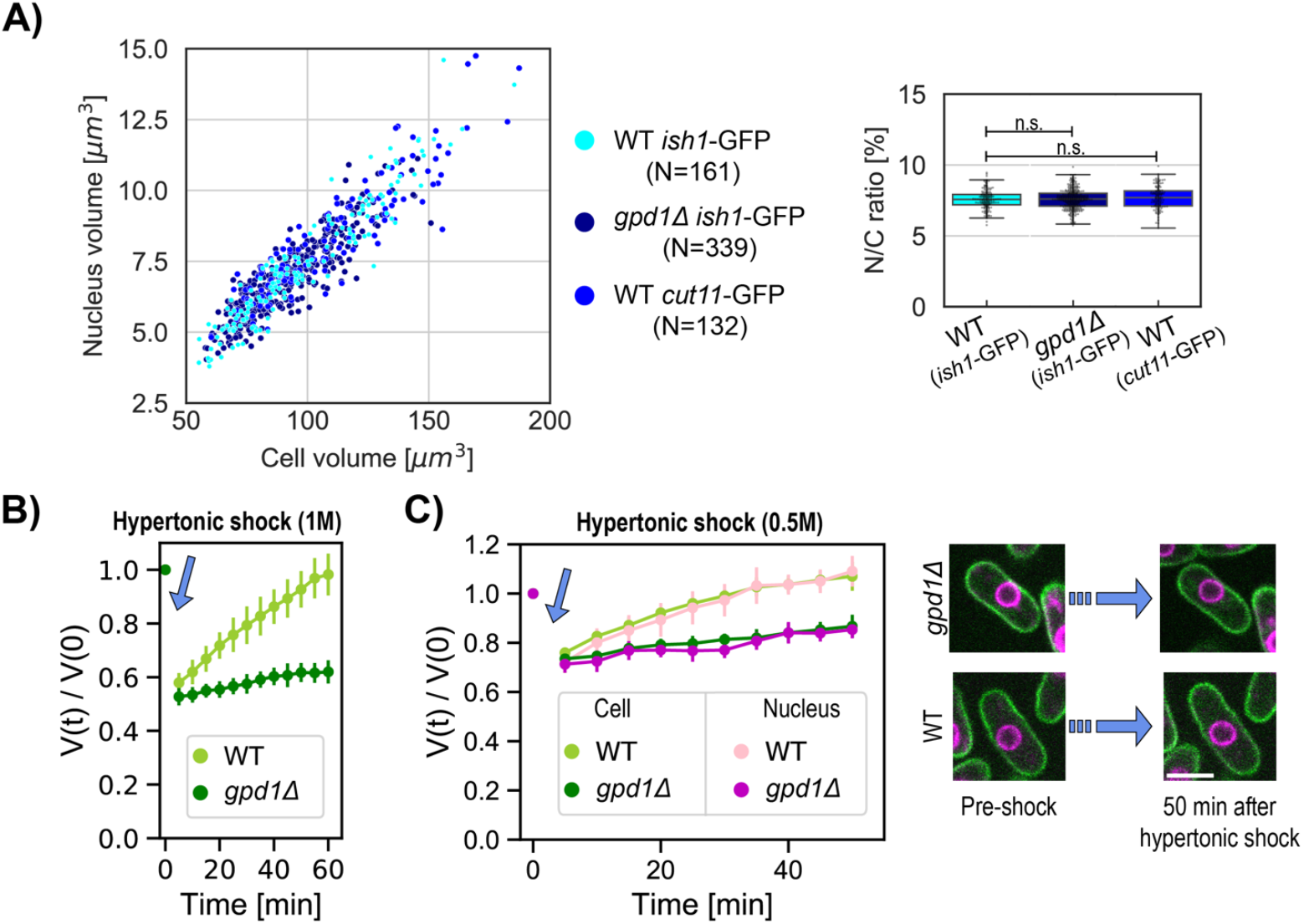
3D image analysis methods and use of an osmotic adaptation mutant allow for robust volume measurements. Previous fission yeast cell studies estimated nuclear and cell volumes from length and single width measurements using assumptions of symmetric ellipsoid or cylindrical geometry (Neumann and Nurse, 2007; Kume *et al*., 2017; Facchetti *et al*., 2019; Lemière, Ren and Berro, 2021). As the shape of fission yeast cells are not perfectly symmetric ellipsoids, we determined volumes using a 3D segmentation approach (Machado, Mercier and Chiaruttini, 2019). To minimize the adaptation responses to osmotic stress, most of these studies were done with cells with a *gpd1*Δ background, which is defective in glycerol synthesis responsible for the rapid volume adaptation to osmotic stresses (Hohmann, 2002). **(A)** N/C ratio is maintained in distinct cellular backgrounds. Scatter plot of cell size and nuclear size for asynchronous cells in growth medium. Right, Box and whisker plots of the N/C ratio for the three strains. Our 3D measurements of mean cell volume (97.5±27.1 μm^3^), nuclear volume (7.3±2.1 μm^3^), and the N/C ratio (7.5±0.8) of a population of asynchronous cells were consistent with previously reported values that used different image analysis methods. (Neumann and Nurse, 2007; Kume *et al*., 2017). For all box and whiskers plots, the horizontal line indicates the median, the box indicates the interquartile range (IQR) of the data set while the whiskers show the rest of the distribution within 1.5*IQR except for points that are defined as outliers. Statistical difference compared with an unpaired t-test. These data show that the N/C ratio measurements was not affected by the *gpd1*Δ background, or by use of different nuclear envelope markers ish1-GFP and cut11-GFP. **(B&C)** Time course of volume adaption in response to hyperosmotic shocks of 1M sorbitol (B) and 0.5M sorbitol (C). Normalized cell volume dynamics WT (chartreuse, N=12 cells) and *gpd1*Δ (green, N=12 cells) after 1 M sorbitol shock, mean ±STD. **(C)** Normalized cell and nucleus volume dynamics after 0.5 M sorbitol shock in WT (N=12) and *gpd1*Δ (N=10) background cells, mean values ±STD. These measurements of volume adaption enabled us to define time windows in which acute volume changes can be measured.

**Figure S2.**
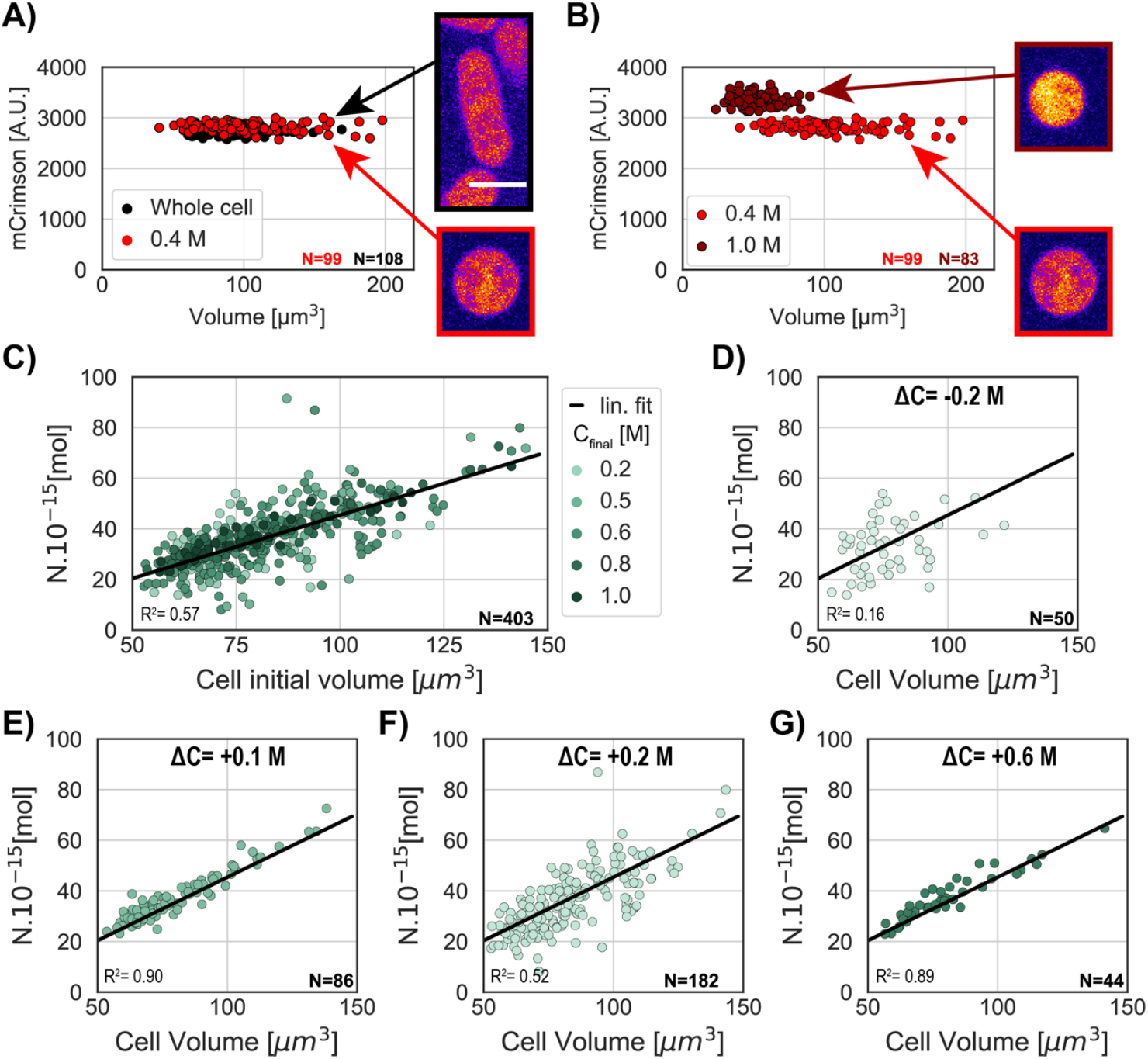
Additional evidence that protoplasts behave as ideal osmometers. **(A)** Defining an isotonic medium for protoplasts. The osmotic pressure of the medium changes the protoplasts’ volume and concentration of proteins due to addition or removal of water from the cell. To assess cytoplasmic concentration, we monitored fluorescence intensity of a protein E2-mCrimson expressed from the ACT1 promoter, which has been shown to normally maintain a largely constant concentration in whole cells throughout the cell cycle (Al-Sady et al., 2016; Knapp et al., 2019). mCrimson fluorescence intensity and cellular volume in a population of whole cells in isotonic medium (black) were similar to those of protoplasts in same medium supplemented with 0.4 M sorbitol (red), demonstrating that this is the isotonic condition for these protoplasts. Right panels, mid focal plane image of a cell (top) and protoplast (bottom) expressing mCrimson. **(B)** In contrast, comparison of mCrimson fluorescence intensities in protoplasts in 1 M sorbitol (dark red) and 0.4 M sorbitol (red), showed that 1M sorbitol led to higher protein concentration than in isotonic conditions. **(C-G)** Number of intracellular osmolytes (N) was measured as described in Methods. Consistent with an ideal osmometer, N is directly proportional to the change in cell volume under osmotic shocks. N is proportional to the cell initial volume and does not depend on the range of osmotic shock used to probe the cells (Methods). R^2^ values are R-squared values for linear regression. Scale bar = 5 μm.

**Figure S3.**
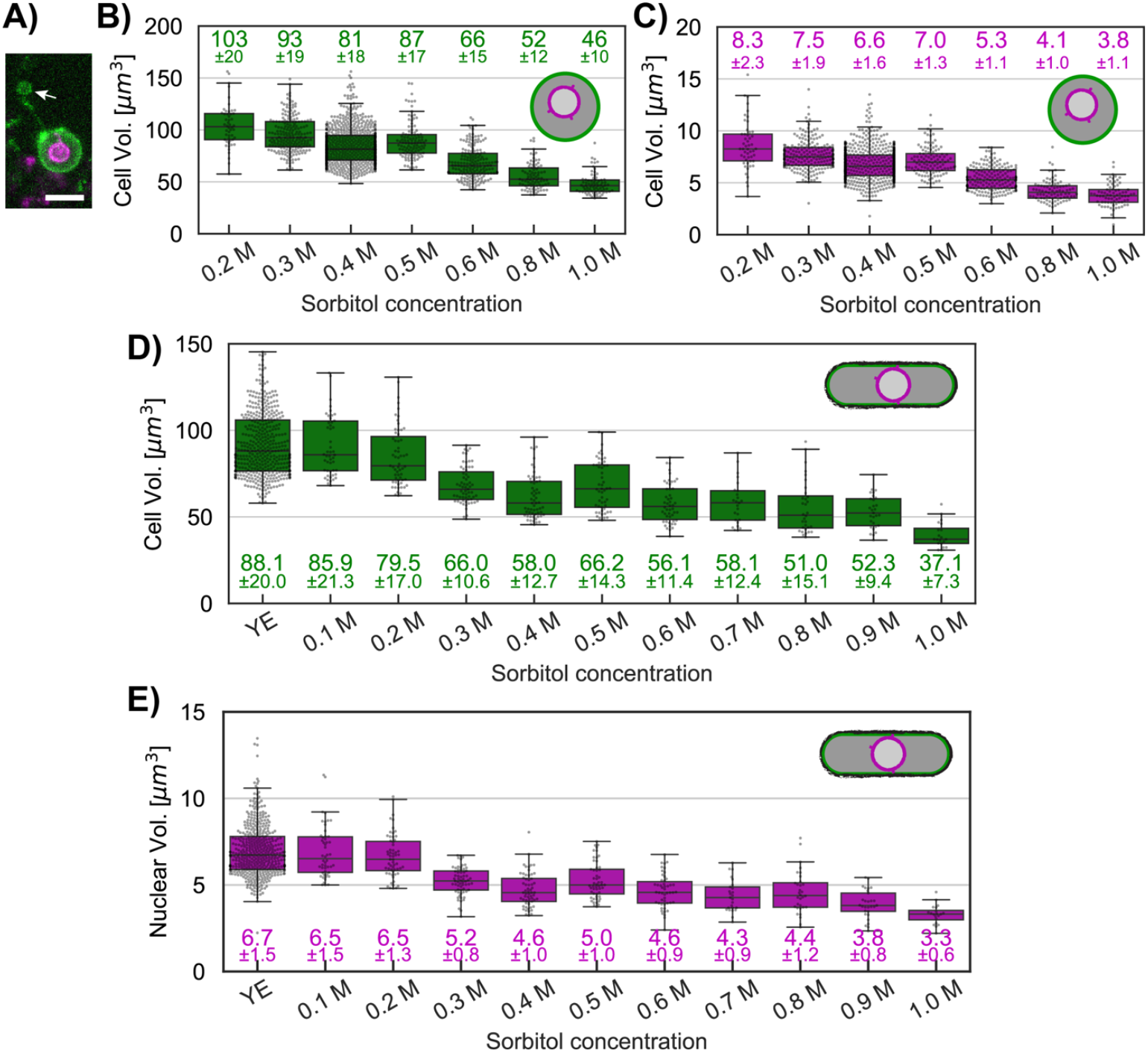
Measurements of cellular and nuclear volumes under osmotic shocks in protoplasts and whole cells. **(A)** During protoplast preparation, a small portion of the cytoplasm was sometimes lost when the protoplast extruded out of the remaining cell wall. Image depicts a sum projection image of a protoplast with a portion of the plasma membrane (green) left behind (arrow). This behavior led to initial measurements that N/C ratio was slightly increased in some protoplasts compared to walled cells (Figure 3B, D). To address this issue, for Figure 3A we therefore focused our analysis on cells that did not show any loss of cytoplasm during protoplasting. **(B)** Box and whisker plots for protoplasts showing individual cells volumes per osmotic condition described in main Figure 3. **(C)** Same as **(B)** for protoplasts nuclei. **(D)** Box and whisker plots for whole cells showing individual cells volumes per osmotic condition described in main Figure 3. Mean ±STD are indicated under each condition. **(D)** Same as **(E)** for whole cells nuclei. For all box and whiskers plots, the horizontal line indicates the median, the box indicates the interquartile range (IQR) of the data set while the whiskers show the rest of the distribution within 1.5*IQR except for points that are defined as outliers.

**Figure S4.**
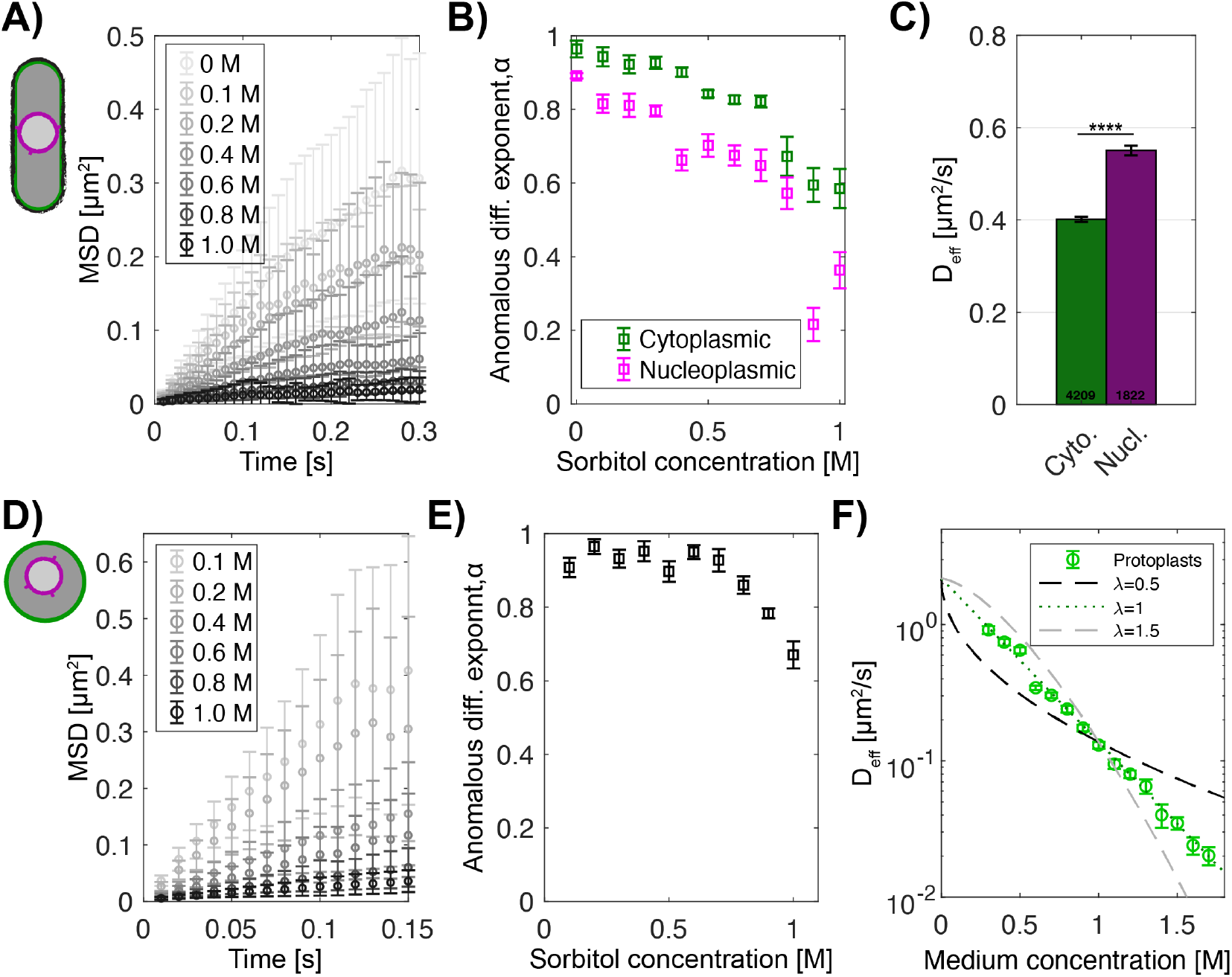
Comparison of the cytoplasmic and nucleoplasmic GEMs diffusion and anomalous exponent under osmotic shocks. **(A)** Example of MSD plots for cytoplasmic GEMs in whole cells under isotonic condition (0 M) or hypertonic shock (0.1-1.0 M of sorbitol). **(B)** Cytoplasmic (green) and nucleoplasmic (purple) GEM anomalous diffusion exponents for whole cells under osmotic shock. **(C)** GEMs effective diffusion coefficient (mean ± SEM) is slower in the cytoplasm (green) than in the nucleoplasm (purple) in whole cells in YE medium. Numbers indicate the number of tracks, p-value<0.0001 Mann-Whitney U test. **(D)** Example of cytoplasmic GEMs MSD plots in protoplasts in isotonic condition (0.4 M) or osmotically shocked (0.1-0.2 M and 0.6-1.0 M). **(E)** Anomalous diffusion exponent *α* obtained from linear fits of the MSD plots presented in (D). **(F)** Effective diffusion coefficient of cytoplasmic GEMs in protoplasts for various medium concentrations. Dashed lines, Phillies’ model for diffusion from three power law values: *λ*=0.5 (black), *λ*=1 (green), and *λ*=1,5 (gray).

**Figure S5.**
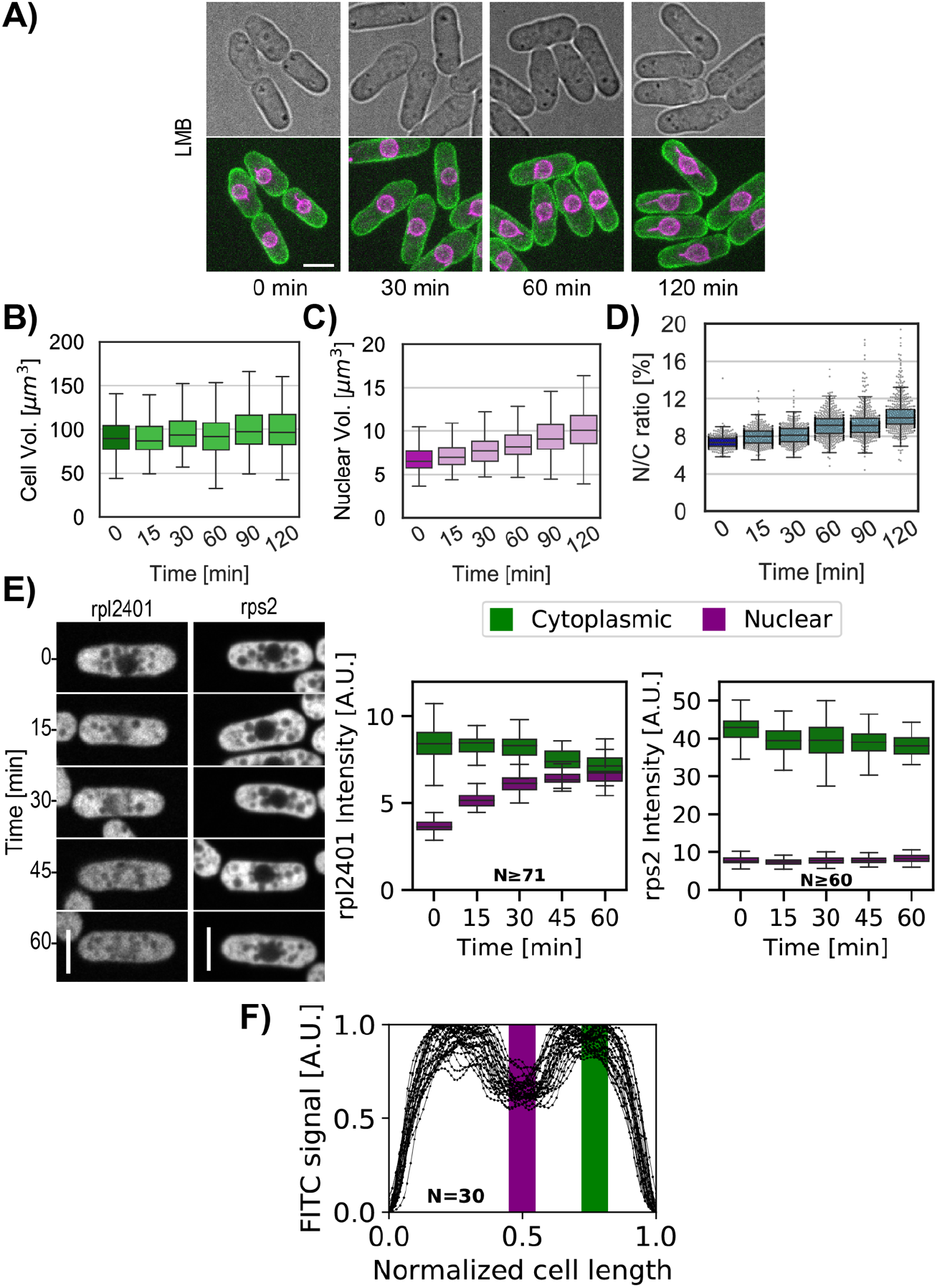
Effects of LMB on the N/C ratio and ribosomal protein localization. **(A)** Bright field mid-focal plane (top) and max Z-projection (bottom) images of the plasma membrane (green) and nuclear membrane (purple) of whole cells treated with LMB over time. Cells were selected to have approximately the same size. **(B-C)** Whole-cell volume and nuclear volume of distinct populations of cells treated with LMB and initial condition. **(D)** N/C ratio of a population of cells treated with LMB (light blue) and initial condition (blue). N≥463 per time point. **(E)** Cells expressing chromosomally tagged proteins that mark the large ribosomal subunit (Rpl2401-GFP) and the small ribosomal subunit (Rps2-GFP) were treated with LMB and imaged over time. Mid focal plane confocal images and quantitation of their relative fluorescence intensities are displayed. **(F)** To reveal the distribution of total protein cells were fixed, treated with RNAse and stained with FITC dye. FITC fluorescence intensities along the normalized cell length were measured. FITC intensities in nuclear and cytoplasmic regions are defined by the signal in purple and green bar. Only a subpopulation of cells was plotted. Scale bar = 5 μm.

**Figure S6.**
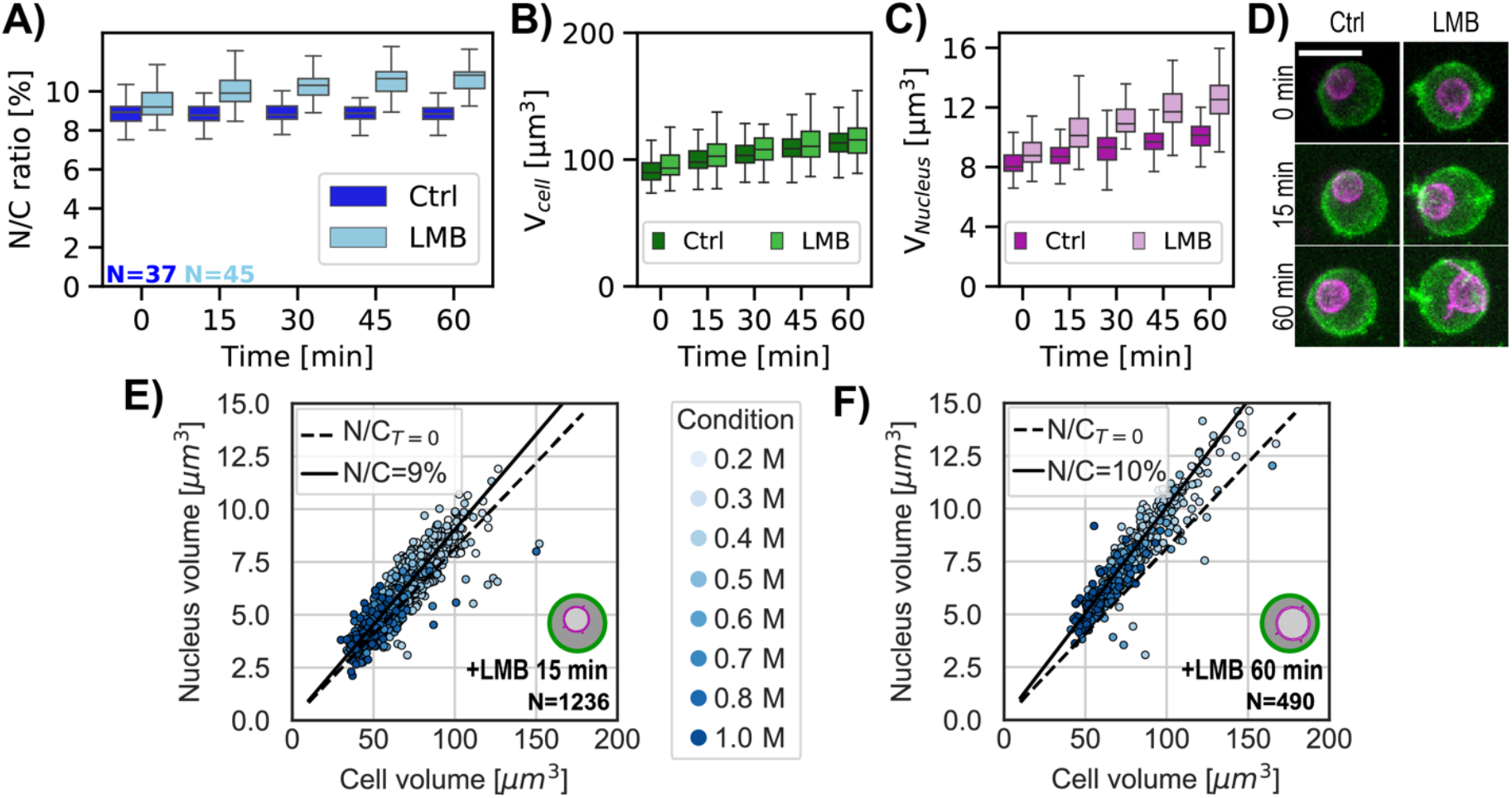
Effects of LMB and osmotic shifts on protoplasts. **A-C)** Time course of N/C ratio cellular volume and nuclear volume in individual control and LMB-treated protoplasts. **(D)** Z-sum projection image of the plasma membrane (green) and nuclear membrane (purple) of protoplasts over time treated with LMB (right) or not (Ctrl, left). **(E)** Scatter plot of cell size and nuclear size for protoplasts in isotonic condition treated with LMB for 15 min (YE + 0.4 M sorbitol) and immediately following osmotic shock. Black line, N/C ratio of cells under isotonic condition (N/C=9%). Dashed line, N/C ratio of cells in isotonic condition before the addition of LMB (N/C_T=0_). **(F)** Same as (E) for protoplasts treated with LMB for 60 min. Scale bar = 5 μm.

**Figure S7.**
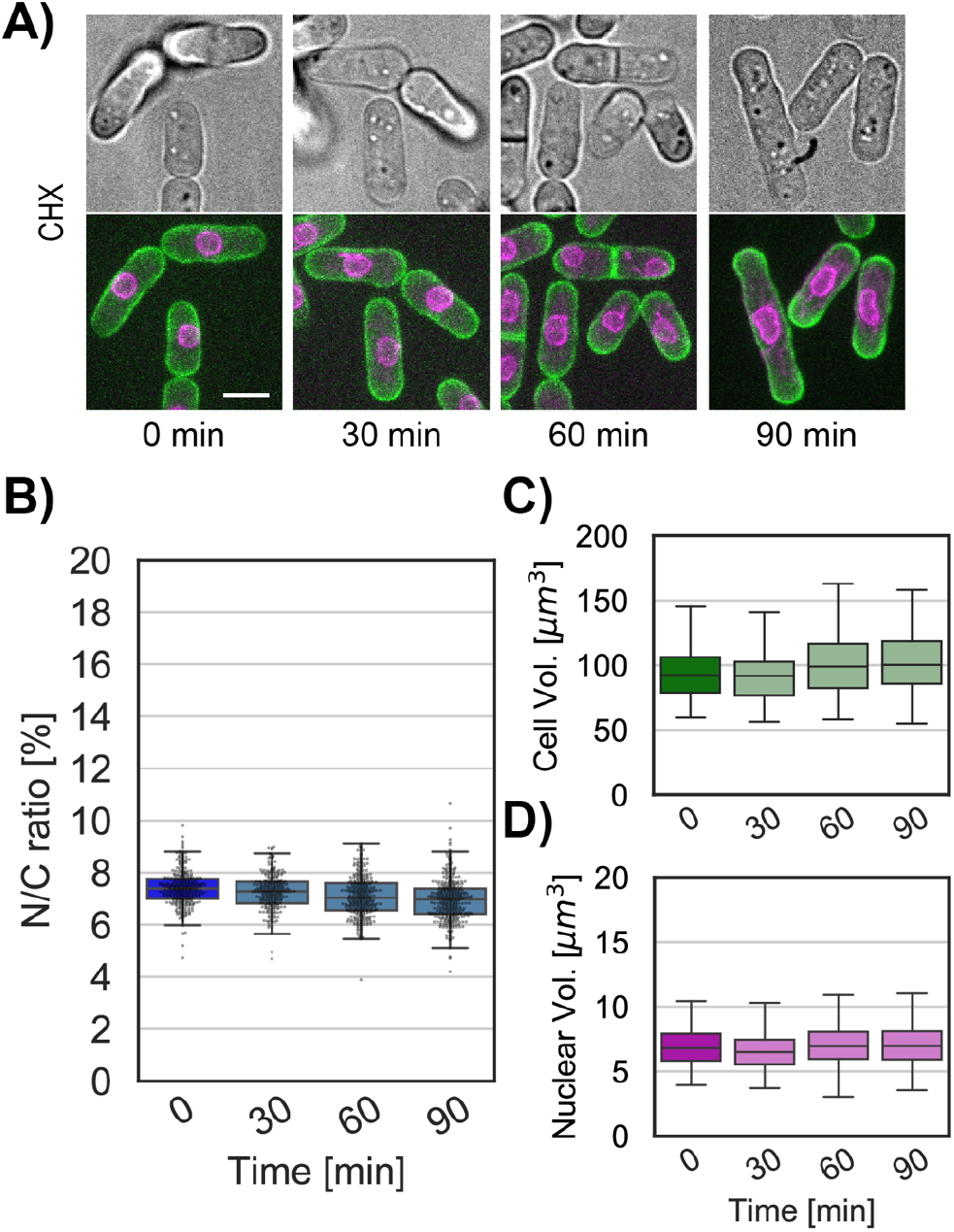
CHX treatment produces no detectable change in N/C ratio. **(A)** Bright field (top) and max Z-projection (bottom) overlay of the plasma membrane (green) and nuclear membrane (purple) of whole cells treated with CHX over time. Cells were selected to have approximately the same size. **(B)** Box and whisker plot of a populations of whole-cell N/C ratio dynamics under treatment with CHX (light blue) and initial condition (blue). **(C-D)** Box and whisker plot of the volume and nuclear volume of single whole cells used to construct (B). N≥253 per time point. Scale bar = 5 μm.

**Figure S8.**
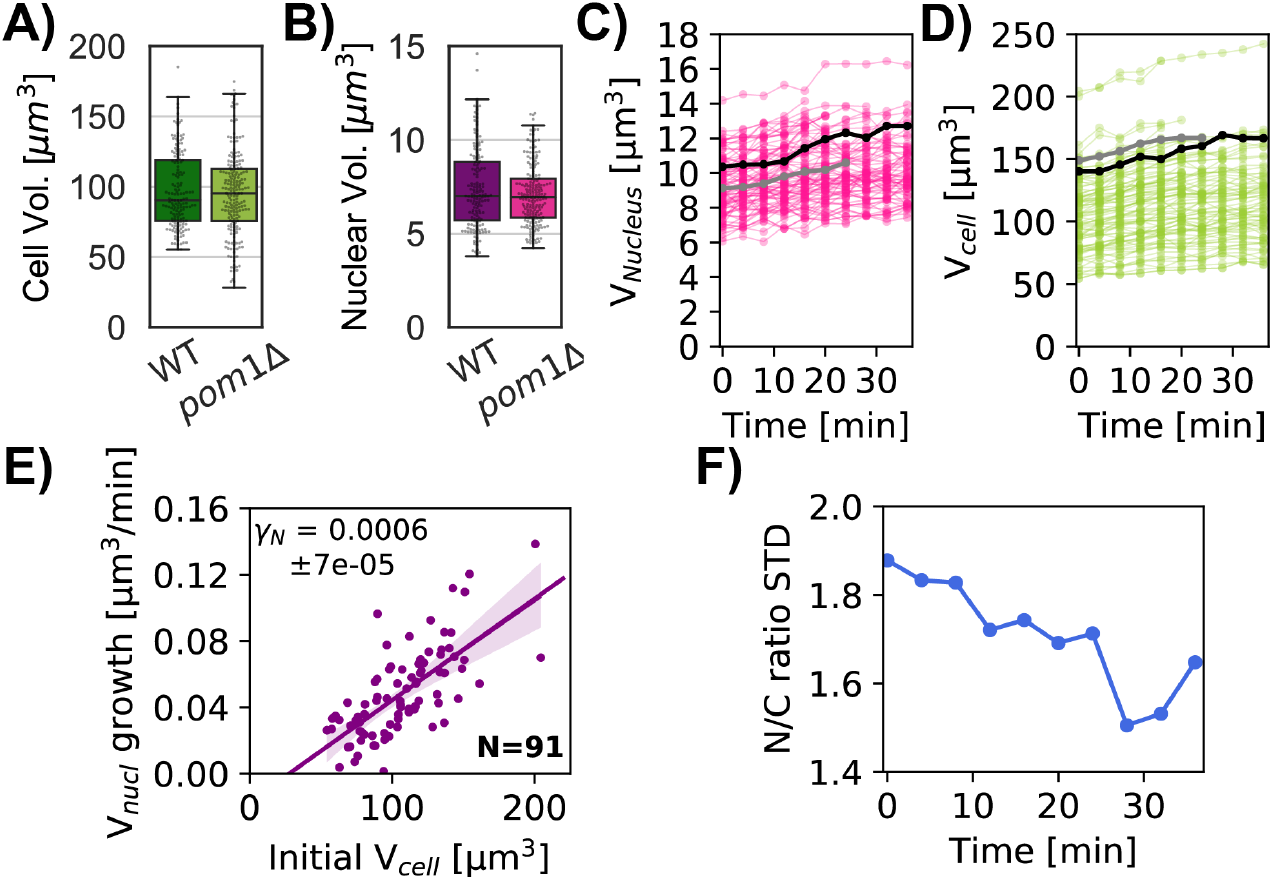
Quantitation of cellular and nuclear growth in *pom1*Δ mutant cells. **(A)** Wildtype and *pom1*Δ whole-cell volumes used for Figure 7A. **(B)** Same as A for nuclear volumes of WT (purple) and *pom1*Δ (pink) cells. **(C)** Nuclear and **(D)** cell volume dynamics for *pom1*Δ cells. Gray and black lines show tracks corresponding to two selected cells. The gray track represents a cell that divided before the end of the experiment. **(E)** Nuclear growth rate as a function of a cell’s initial volume. Linear regression is shown in purple with the slope *γ*_N_. **F**) The standard deviation (STD) of the N/C ratio measured for 91 *pom1*Δ cells decreases over time.

## Appendix

## A Mathematical Model

We describe a simple theory of nuclear size control based on osmosis, in which the balance of solute particles in the cytoplasm and nucleoplasm determines a unique steady-state nucleus to cell (N/C) ratio.

Our theoretical model is based on the physical mechanism of osmosis. For a perfect osmometer, Van’t Hoff’s law asserts that the osmotic pressure on a body is determined solely by the relative concentrations of solute particles inside and outside of the body [1]. In the context of nuclear size control, we idealize the cell and nucleus as two nested osmometers—with the cell containing the nucleus—and interpret van’t Hoff’s law in terms of the concentrations in the cytoplasm, nucleoplasm, and extracellular space.

Cells are densely packed with ions, proteins, and other biomolecules. Without additional constraints, the cell would simply swell osmotically until its internal concentration reached that of the surrounding medium. The elastic cell membrane opposes this osmotic pressure by membrane tension. In this note, we describe how one may calculate steady-state sizes predicted by the balance of forces between membrane tension and osmotic pressure via van’t Hoff’s law. The steady-state size depends on the cell’s elastic properties along with the solute concentration (or number of molecules).

Our model is based on previous investigations in the case of a single compartment, e.g. vesicles consisting of a single lipid bilayer membrane, which were validated by hydrodynamic simulations [2]. In this previous study, the steady-state size could be calculated based on two parameters: the surface tension of the membrane and the number of solute molecules.

Following [2], we assume that at steady-state each membrane is in a state of mechanical equilibrium. This implies that osmotic pressures *P*_osm_ are balanced by mechanical tension *T* in each membrane. The osmotic pressure *P*_osm_ satisfies van’t Hoff’s law, which states that *P*_osm_ = *ck_B_T* in which *c* is the concentration of solute molecules, *k_B_* is Boltzmann’s constant, and *T* is the temperature. Whereas in [2] the assumption of mechanical equilibrium is imposed across a single spherical membrane, here we apply this assumption across both the inner and outer membranes, which results in the coupled equations

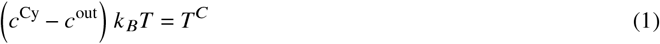

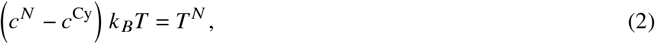

where *T^C^* and *T^N^* are the tensions on the cell and nuclear membranes, respectively, and the external concentration *c*^out^ is a parameter that may be tuned experimentally.

In the case of a spherical membrane of radius *R* subject to surface tension *σ*, Young-Laplace’s law gives *T* = 2*σ*/*R*. Assuming the inner and outer membranes have the same surface tension *σ*, eqs. (1) and (2) becomes

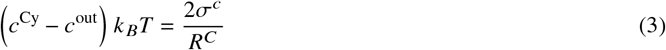

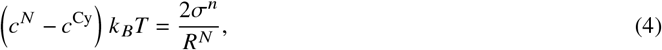

in which *R^C^* and *R^N^* are the radii of the cell and nucleus and *σ^C^* and *σ^N^* are the respective membrane tensions. Assuming the nucleus and cell are both spherical yields Eqs. (1) and (2) of the Main Text upon expressing *R^N^* and *R^C^* in terms of their respective volumes.

### A.1 Prescribed concentration

It follows immediately from (3) that to obtain a finite size nuclear membrane the solute concentrations must satisfy *c^N^* > *c^Cy^*, in which case the nuclear membrane radius is

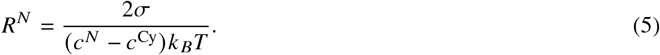

If *c^N^* – *c*^Cy^ ≤ 0, both surface tension and osmotic pressure will exert inward forces on the nuclear membrane and its radius will shrink to zero.

Similarly, if the cytoplasm has a prescribed concentration *c*^Cy^ > *c*^out^, the resulting steady-state radius *R^c^* will satisfy

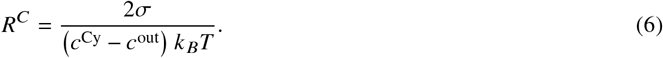

As above, if *c*^Cy^ ≤ *c*^out^ both surface tension and osmotic pressure will exert inward forces on the cell membrane and its radius will shrink to zero. In the case of prescribed concentrations *c^N^* > *c*^Cy^ > *c*^out^, the resulting ratio of nuclear to cell radii is therefore

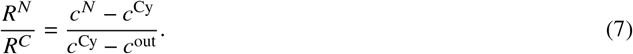

### A.2 Prescribed solute number

Alternatively, rather than specifying the concentrations, we may instead specify the total numbers of solute molecules *N^N^* and *N^C^* contained in the nucleus and cell (so that *N^C^* − *N^N^* is the number contained within the cytoplasm, see Fig. 1(A) in the Main Text). In this case, van’t Hoff’s law may be written in terms of the number of solute molecules *N* and volume 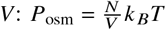. This results in two coupled equations for the membrane radii:

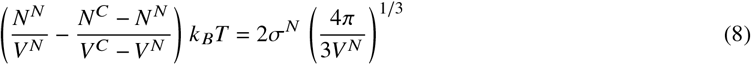

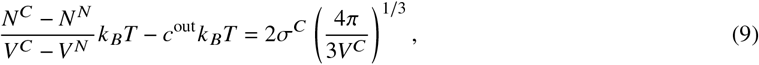

where we have used the relationship *V* = (4/3) *πR*^3^ for the volume of a sphere to calculate the radii used in Young-Laplace’s law. We remark on the importance of accounting properly for the membrane permeability in the osmotic balance. That is, only those solutes which are impermeable to the membrane may contribute osmotic pressures. In our model, we assume that the cell membrane is impermeable to both macromolecules and ions, i.e. that ion leakage and pumping are not significant on the timescales of interest, whereas the nucleus is impermeable to macromolecules but not to ions. This is why *N^C^* must include the number of ions in (8) but not in (9).

In the case of zero surface tension (*σ^N^* = *σ^C^* =0), we may simplify the equations above. In this case, (8) simply states that the concentrations are equal in the two compartments, i.e.

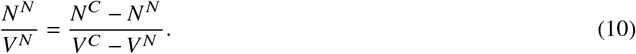

It follows that the total concentration satisfies *N^C^*/*V^C^* = *N^N^*/*V^N^*, so that the N/C ratio is given by

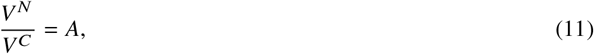

where *A* = *N^N^*/*N^C^* is the ratio of solute molecules in the nucleus and cell. Moreover, from (8) and (9), we have

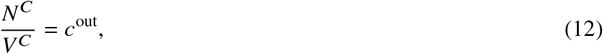

so that

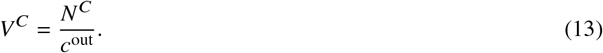

Assuming the total number of solute molecules are fixed, the above relation (13) predicts a linear relationship between the cell volume and inverse external concentration, with a *y*-intercept at *V^C^* = 0.

Experiments including Figs. 2(D)-(E) in the Main Text agree with the linear correlation but reveal a nonzero y-intercept. To account for the nonzero y-intercept in the data, we introduce a non-osmotic volume *b* in both the cell and nucleus, so that the osmotic pressure has the modified form 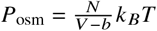. This results in the following modifications to eqs. (8) and (9):

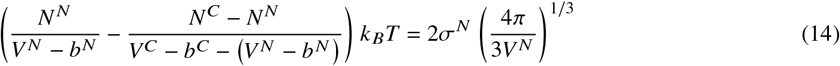

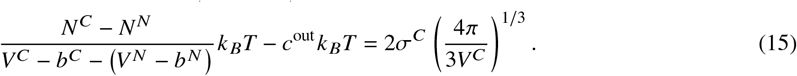

Note that *b^C^* – *b^N^* is simply the cytoplasmic non-osmotic volume.

Upon including the non-osmotic volume in this manner, it is straightforward to show that for the case of zero surface tension (*σ*^N^ = *σ^C^* = 0) the analogues of (11) and (13) are

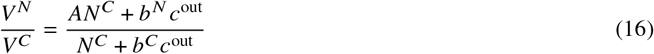

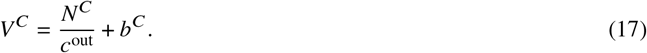

Some special cases of (16) are especially revealing. In the case *b^N^*/*b^C^* = *A* we have *V^N^*/*V^C^* = *A* for all *c*^out^, recovering the result (11) obtained without non-osmotic volume. On the other hand, in the limit *c*^out^ → ∞, we have *V^N^*/*V^C^* → *b^N^*/*b^C^* independent of *A*. Although here we have shown this result only in the case of zero nuclear and cell membrane tension, it holds more generally whenever the nuclear membrane tension is zero, as shown in D.

Note further that, according to (17), in the case *σ^N^* = *σ^C^* = 0 the cell size *V^C^* is determined only by the total number of solute molecules *N^C^* and is independent of the relative ratio *N^N^*/*N^C^*. In practice, we find that the cell size stays nearly constant over typical non-zero values of *σ^N^* as well, whereas the nuclear volume and N/C ratio increase as the relative nuclear abundance *N^N^*/*N^C^* increases.

In the case of non-zero surface tension, the equations are more difficult to solve analytically. However, it is straightforward to solve eqs. (14) and (15) numerically over a range of external osmolarities *c*^out^ and to do so we use the command fsolve in Matlab.

**Table A1:**
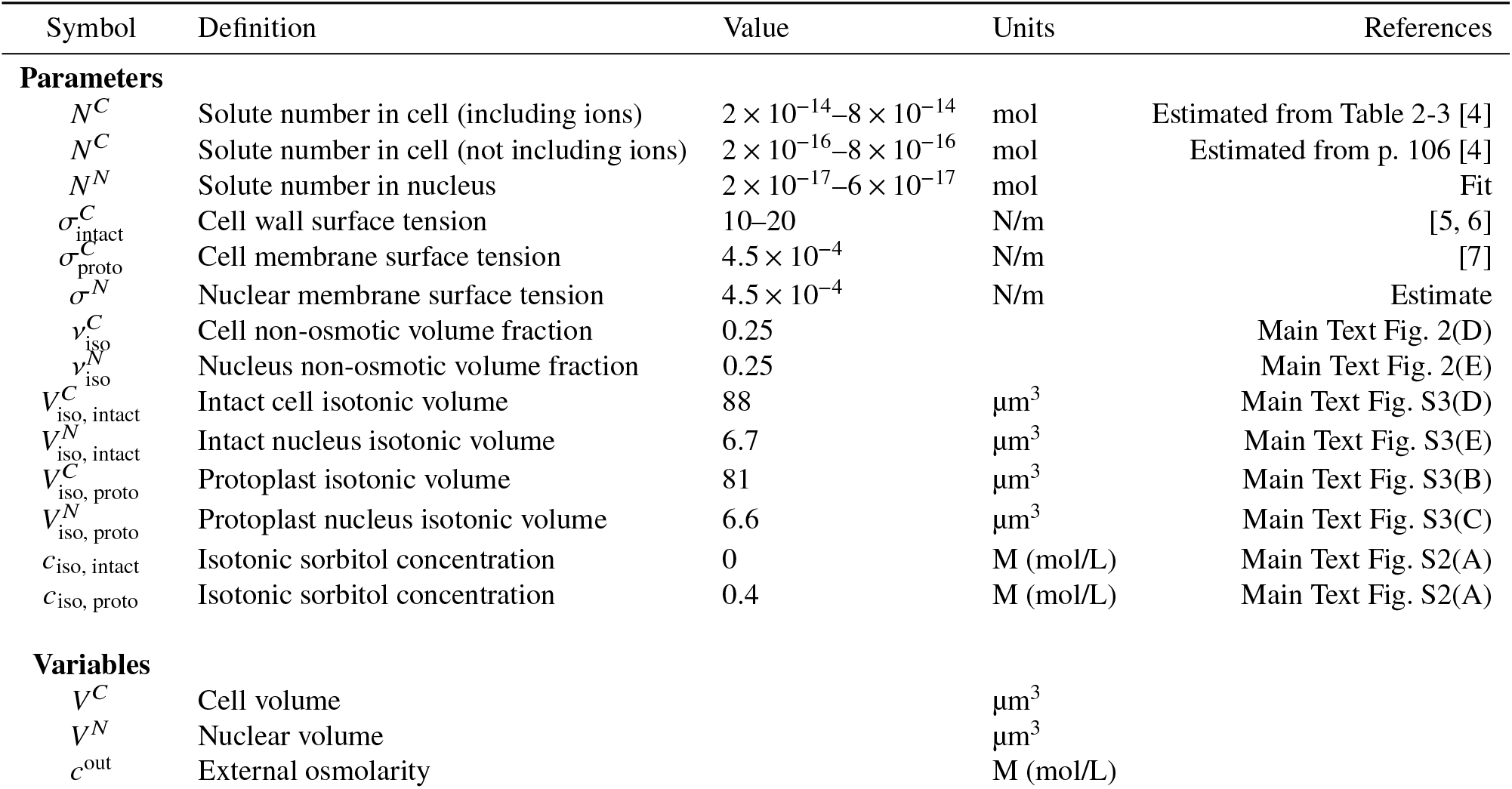
Parameter values and definitions

### A.3 Interpretation of solute molecules

An important question is whether to include the number of macromolecules, ions, or both in variables *N^N^* and *N^C^*. This requires careful consideration of the nature of the compartment and the molecules to which its boundary is permeable.

We therefore consider the relevant osmolytes at both the nuclear and cell membranes. The nuclear pore complex is permeable to ions which can freely cross the nuclear membrane [3]. Therefore only macromolecules are able to sustain a concentration gradient, and *N^N^* and *N^C^* in the force balance equation (repeated here for convenience) represent the numbers of macromolecules only (not including ions):

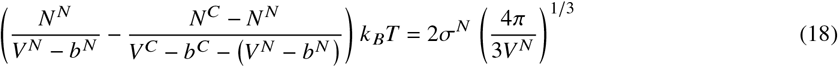

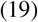

At the cell membrane, on the other hand, there are ion pumps that are able to maintain a concentration gradient. Therefore we must include two general classes of osmolytes: macromolecules and ions. In the case *σ^N^* = 0 explored above, this leads to the following force balance at the cell membrane

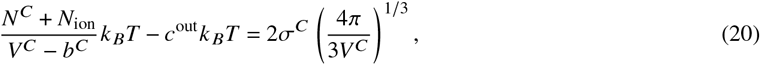

where *N^C^* and *N*_ion_ represent the numbers of macromolecules and ions inside the cell, respectively. In effect, the overall size set by (20) is determined by *both* the macromolecules and the ions, whereas only the number of macromolecules and not the number of ions is important for the N/C ratio determined through (18).

It is important to note that the number of ions contained in the cell is greater than the number of proteins by approximately two orders of magnitude. This effectively decouples the two equations (14) and (15), since the cell size is dominated by the number of ions, whereas the N/C ratio is dominated by the numbers of proteins.

## B Model results

### B.1 Response to increasing solute content

Inspired by experiments in which nuclear export is blocked using the drug Leptomycin B (LMB), we next consider the effect of moving solute molecules from the cytosol to the nucleus on the N/C ratio. We start with the following expression for the N/C ratio in the case *σ^N^* = *σ^C^* = 0, in which case:

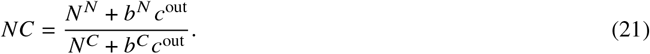

To determine the effect of increasing the number of nuclear solute molecules *N^N^* while holding the total number of solute molecules *N^C^* fixed, we compute

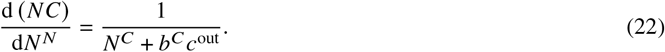

According to (22), upon importing a (small) number Δ*N* of solute molecules into the nucleus, we expect a corresponding change of 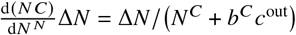 in the N/C ratio. If we enter the estimated values of parameters from Tab. A1, this implies that approximately 10^−18^ mols solute must be imported into the nucleus to generate a 1% change in the N/C ratio. The model predicts that importing a single solute molecule changes the N/C ratio by 2 × 10^−7^%.

Moreover, we may use the measured nuclear and cell sizes during LMB experiments to estimate the solute concentrations and numbers of over time. We find that whereas the concentrations are relatively stable for both protoplasts and intact cells during compared to controls, LMB yields an elevated nuclear solute number upon fitting consistent with the experimental trend (Fig. B1).

**Figure B1:**
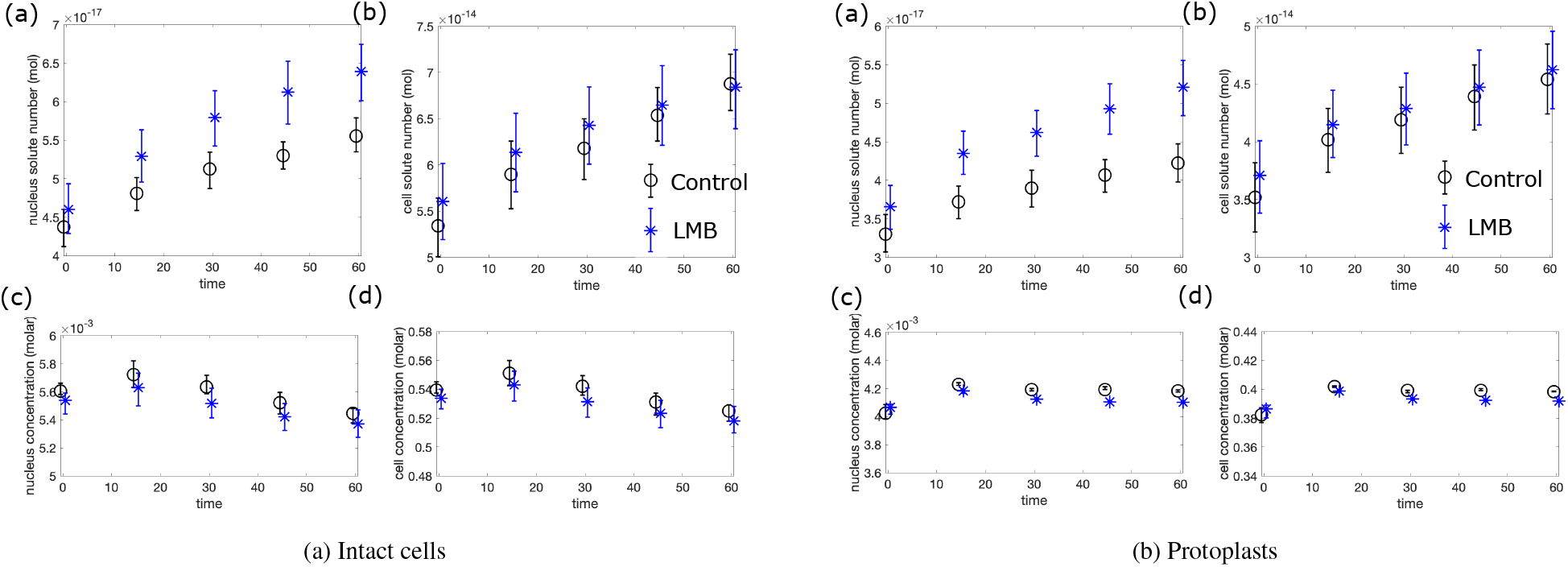
Left panel: intact cells, right panel protoplasts. (a) Nuclear solute number (mol), (b) cell solute number (mol), (c) nuclear concentration (M), (d) cell concentration (M). The model predicts the increase in nuclear solute number in LMB experiments upon fitting to the measured cell and nuclear volumes.

### B.2 Homeostasis through growth

As discussed in Main Text Methods section “Modeling nuclear growth and N/C ratio homeostasis”, the model may be extended to account for cell growth. Upon incorporating cell growth (as discussed next in detail) the model predicts homeostasic behavior in which perturbations in the N/C ratio are corrected over time. Incorporating cell growth in the model leads to two interesting insights. First, exponential growth provides a cell cycle-independent, passive feedback mechanism for correcting aberrations in the N/C ratio. Second, the value of the N/C ratio is determined by the fraction of synthesized proteins that enter the nucleus. These insights provide a plausible genetic explanation for the value of the N/C ratio (the fraction of proteins genetically tagged to enter the nucleus) and raises the question of whether the N/C ratio may be a spandrel, i.e. a simple consequence of this number rather than a quantity controlled by active feedback.

Next, we derive Eq. (4) in the Main Text. Assuming exponential cell growth yields:

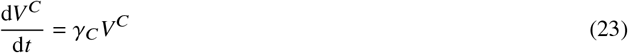

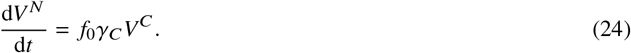

Applying the quotient rule to 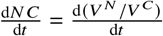 yields

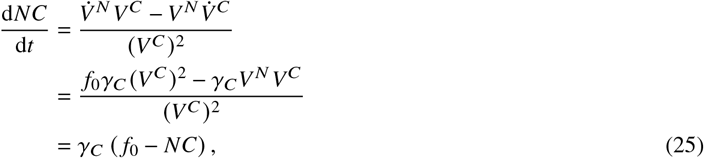

where we have used eqs. (23) and (24) to obtain the second equality. This indicates a passive feedback mechanism through growth that corrects perturbations in the N/C ratio to the steady-state value of *f*_0_.

**Figure B2:**
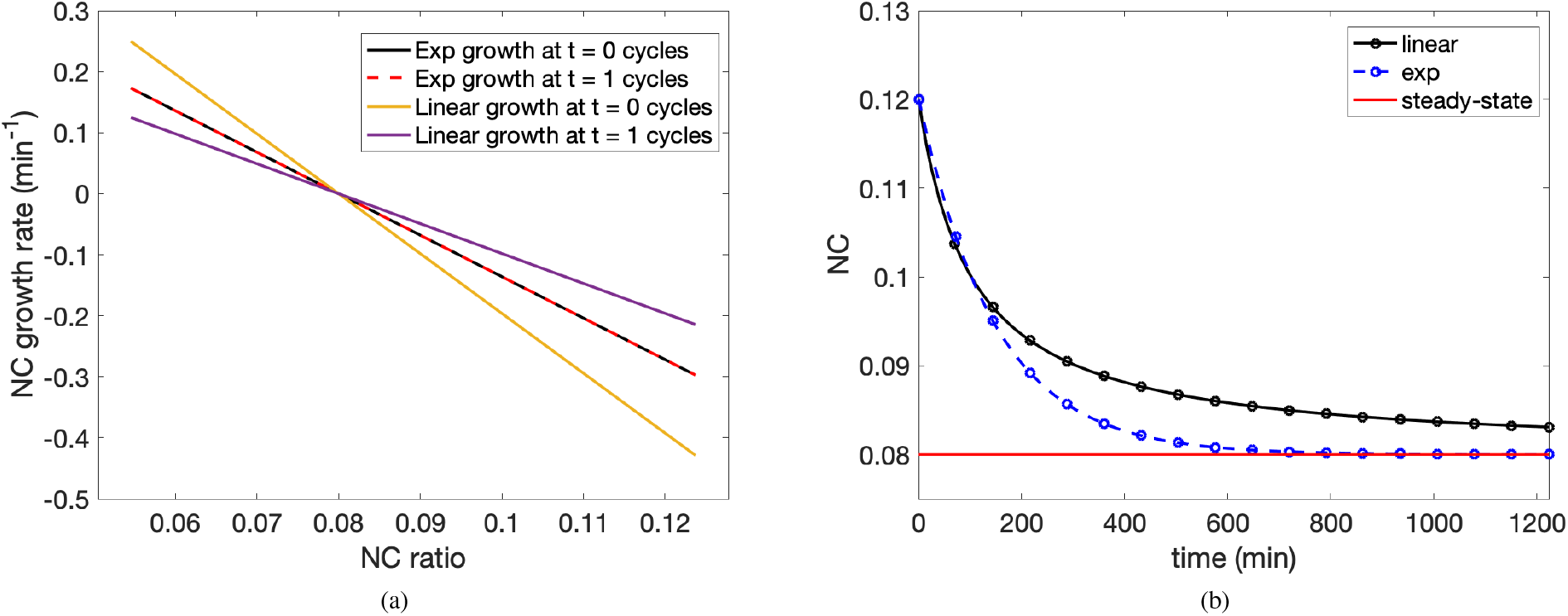
Homeostatic behavior predicted by the model in the case of linear and exponential cell growth. (a) Exponential growth yields a cell cycle-independent homeostasis, (b) Exponential growth leads to a faster recovery of the steady-state N/C ratio in the case of mutants that are unable to divide.

For comparison, we also consider the case of linear growth with a constant growth rate *α*:

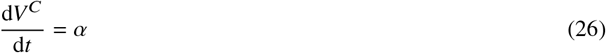

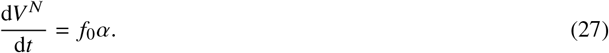

In this case, following the same steps as above yields

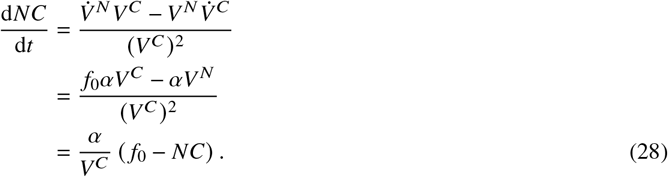

Linear growth also yields a passive homeostasis in the N/C ratio, but with the notable difference that in this case the correction rate is cell-size dependent. This is because of the factor *V^C^* in the denominator of (28). If one calibrates the linear growth model so that the cell doubling time is equal to that of exponential growth, this cell-size dependence manifests as a faster correction rate early on in the cell cycle, and a slower correction rate later on in the cell cycle (Fig. B2(a)).

One consequence of this comparison between homeostatic behavior for linear and exponential growth is that, in mutants in which the cell and nucleus are unable to divide, exponential growth leads to a much more rapid return to homeostasis than linear growth (Fig. B2(b)).

## C Nondimensionalization

We next introduce dimensionless variables in order to consider the limiting cases of large and small surface tension on the inner and outer membranes. In what follows, we modify the notation to make the presentation more general. The inner (e.g. nuclear) variables are denoted by the subscript 1 so that *V*_1_ ≡ *V^N^*, *N*_1_ ≡ *N^N^*, and *σ*_1_ ≡ *σ^N^*, and outer (e.g. cell) variables are denoted by the subscript 2 so that *V*_2_ ≡ *V^C^*, *N*_2_ ≡ *N^C^*, and *σ*_2_ ≡ *σ^C^*. We use co to denote *c*^out^.

Let *Ṽ* = *N*_2_/*c*_0_ denote the characteristic cell volume, *A* = *N*_1_/*N*_2_ be the ratio of solute molecules as above, and 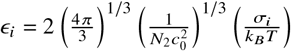 be the nondimensional tension for *i* = 1, 2. Note that *ε*_i_ is indeed a dimensionless group, since surface tension having units of J/m^2^ implies that *σ_i_*/*k_B_T* has units of 1/m^2^, whereas 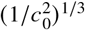 has units of m^2^.

In terms of the dimensionless volumes *v_i_* = *V_i_*/*Ṽ* and non-osmotic volumes *v_i_* = *b_i_*/*Ṽ*, the model eqs. (14) and (15) may be rewritten as

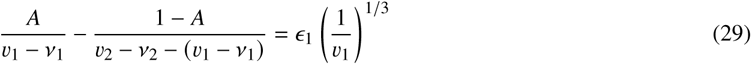

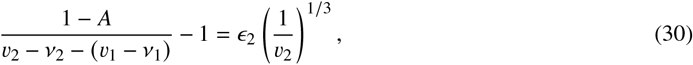

To study the experimentally-relevant case in yeast, we set *ε*_1_ = 0. Therefore

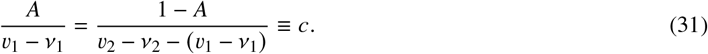

Of course, the overall concentration 1/(*v*_2_ – *v*_2_) must also be equal to *c* as well. This can be shown as follows:

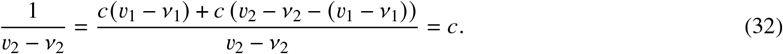

Therefore, in the case of interest *ε*_1_ ≈ 0 we may rewrite (30) as

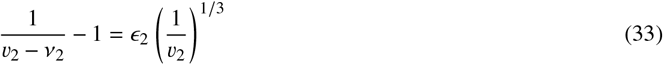

to obtain a single equation for the single unknown *v*_2_. Next, we consider the asymptotics of solutions to this equation in the limits of large and small *ε*_2_.

### C.1 Limit of *ε*_2_ → 0

First, we define an auxiliary variable 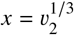 to transform (33) into the following equation in *x*:

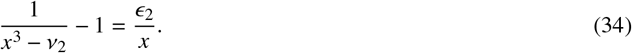

Multiplying through by the common denominator *x*(*x*^3^ – *v*_2_), (34) may be rewritten as a quartic polynomial:

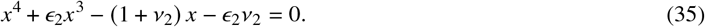

i.e. *f*(*x*) =0 with *f*(*x*) = *x*^4^ + *ε*_2_*x*^3^ – (1 + *v*_2_) *x* – *ε*_2_*v*_2_. Because it a fourth-order polynomial, *f* has four roots, some of which may be complex-valued. Next, we apply the method of dominant balance to approximate the roots. It is straightforward to show that there are two valid balances in the limit *ε*_2_ → 0. First, there is the one obtained by dropping the term involving the small parameter, i.e.

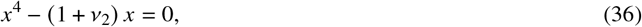

which has a single real root at 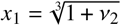 in addition to two complex-valued roots. Another valid dominant balance is obtained by dropping the two higher-order terms:

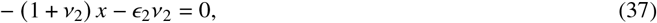

which yields the (non-physical) negative solution *x*_2_ = −*ε*_2_*v*_2_/(1 + *ε*_2_). In practice, we therefore expect solutions to (35) to approach the positive *x*_1_ as the outer surface tension becomes smaller and smaller.

### C.2 Limit of *ε*_2_ → ∞

In this limit, there are two possible balances, each of which retain the large parameter. It is straightforward to show that these are

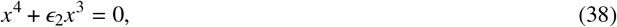

which yields the solution *x*_3_ = −*ε*_2_ and

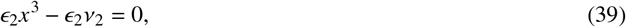

which yields the solution 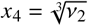. In Fig. C1, we fix *v*_2_ = 0.25 and illustrate the convergence of the roots of (35) to the asymptotic limits given by *x*_1_ and *x*_2_ in the small-*ε*_2_ regime and to *x*_3_ and *x*_4_ in the large-*ε*_2_ regime.

**Figure C1:**
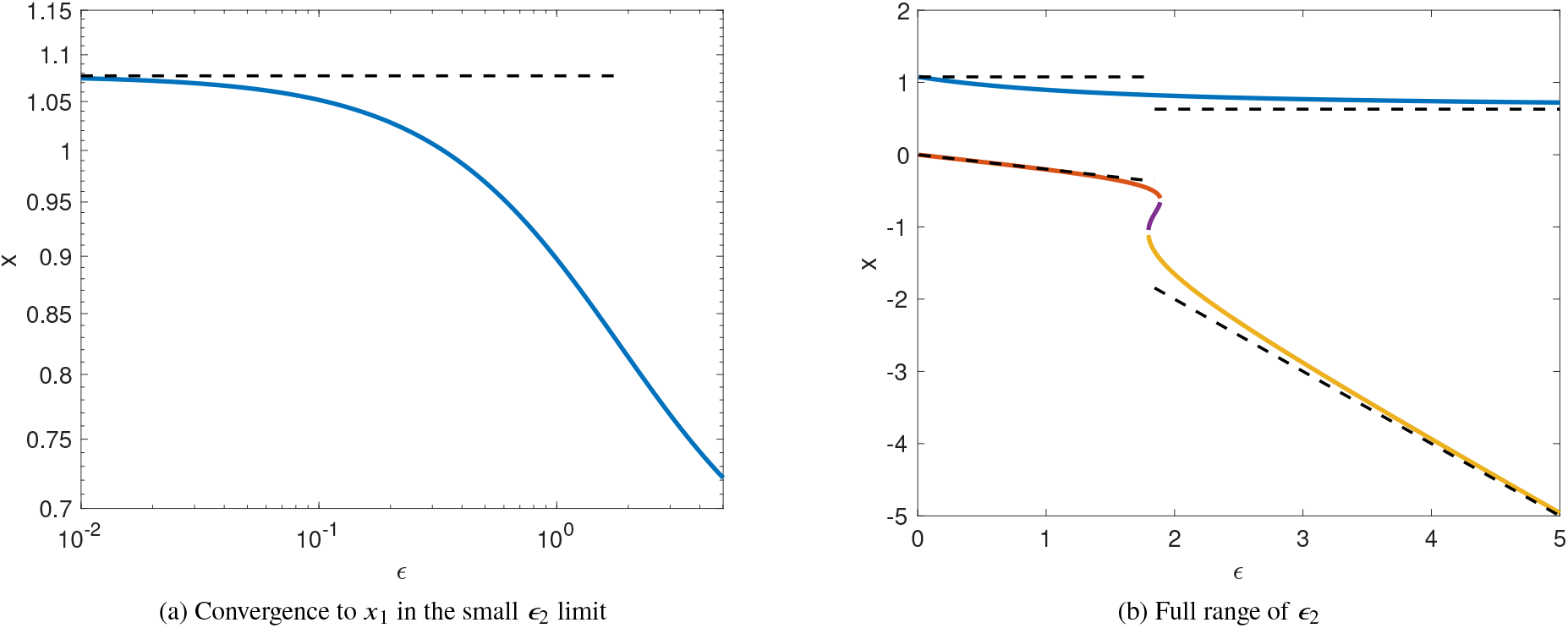
Limiting behaviors with *ε*_2_ in the special case *v*_2_ = 0.25.

### C.3 Transition between limits

As shown in Fig. C1, there is a critical interval in which (35) possesses four real roots. This critical interval is located between the small and large-*ε*_2_ regimes, and therefore we interpret the midpoint of this interval 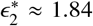 to be the transition between the external osmolarity and surface tension-dominated regimes.

**Figure C2:**
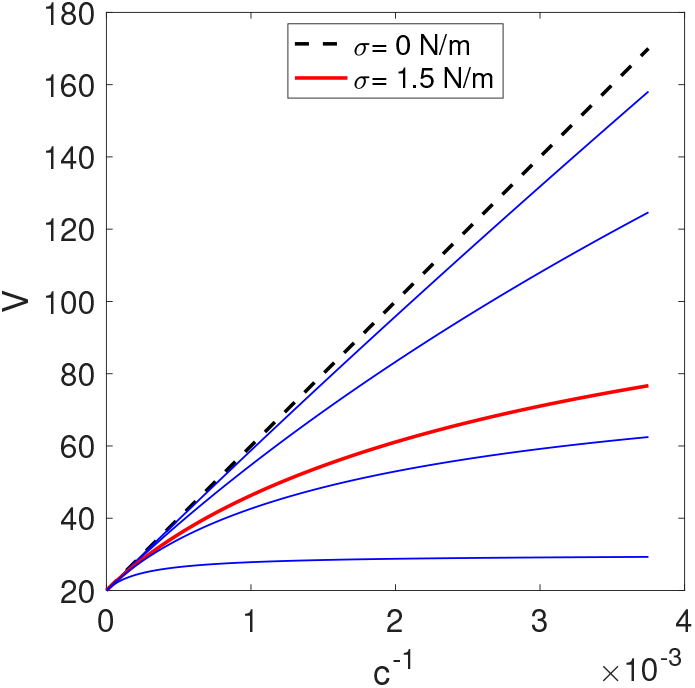
Boyle van’t Hoff plots for different values of the surface tension *σ*_2_. The curve in red corresponds to the critical tension 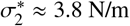, whereas the dotted line corresponds to *σ*_2_ = 0.

To determine the corresponding critical surface tension 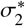, we use the definition of *ε*_2_ together with the values found in Tab. A1 to estimate 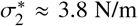. Fig .C2 shows (in red) the Boyle-van’t Hoff plot associated with the critical surface tension 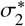, which as expected occurs between the linear (external osmolarity dominated) and highly nonlinear (surface tension dominated) regimes. The fact that 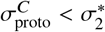 whereas 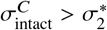 is consistent with the experimental observation that the protoplasts follow an Boyle van’t-Hoff law whereas the intact cell does not.

### C.4 Behavior for large inner tension *σ*_1_

In addition to the limiting case above in which the inner membrane tension *σ*_1_ is negligible, we also consider the solutions to eqs. (14) and (15) in the case of large *σ*_1_ and negligible outer tension *σ*_2_. This situation is likely to be the relevant one for mammalian cells, which do not have a cell wall to provide rigidity to the outer membrane but contain lamins in the nuclear membrane that provide additional structural rigidity compared to yeast.

In Fig. C3, we plot the N/C ratio in this case 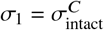 for several choices of the outer tension *σ*_2_. Unlike the inverted limit in which the inner tension is small, here the N/C ratio can be seen to vary considerably in response to osmotic shocks. Based on these model predictions we therefore expect that, unlike the nearly constant N/C ratio maintained by force balance in yeast, maintaining a stable N/C ratio in mammalian cells in response to osmotic shocks would require additional control mechanisms over solute concentrations and/or plasma volumes.

## D Special case: proportional non-osmotic volumes at reference concentration

Here we show that, if the nuclear membrane satisfies *σ^N^* = 0 and the non-osmotic volumes *b^N^* and *b^C^* satisfy 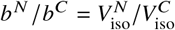, i.e. the non-osmotic volumes are in proportion to the nuclear and cell volumes at some reference concentration, then the N/C ratio satisfies

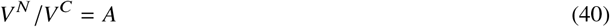

independent of the external concentration *c*^out^.

We proceed with the derivation of (40). In what follows, we use the notation of Section C in which nuclear variables are denoted by the subscript 1 and cell variables by the subscript 2, so that e.g. *V*_1_ ≡ *V^N^* and *V*_2_ ≡ *V^C^*, and in which the

**Figure C3:**
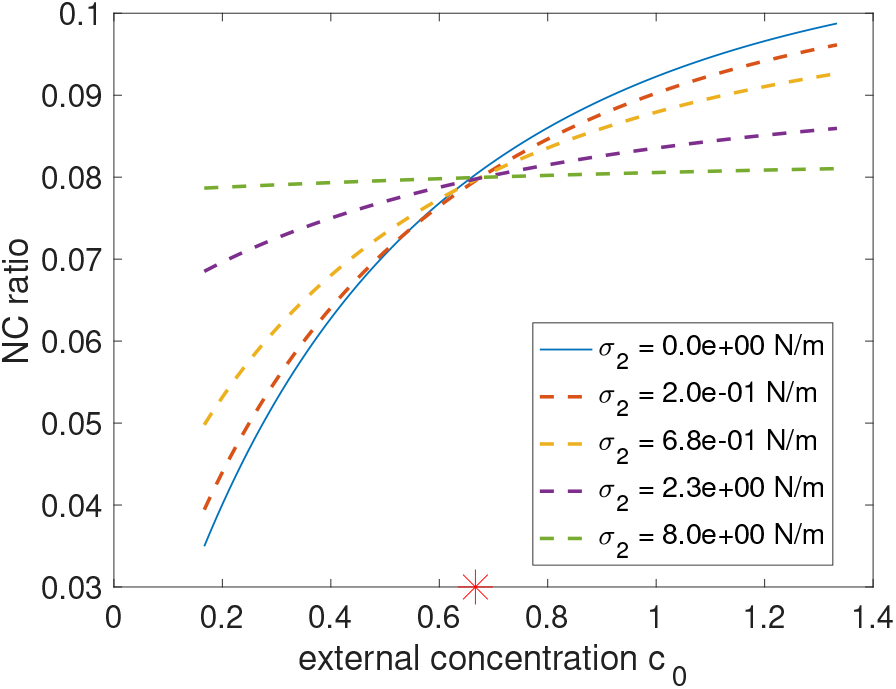
Limiting behavior varying the outer tension *σ*_2_ in the limit of large inner tension 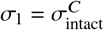.

external concentration is denoted *c*_0_ ≡ *c*^out^. In the tension-free case, the force balance condition analogous to (10) in the presence of non-osmotic volumes is

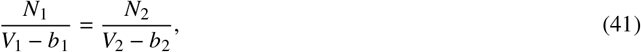

where as above *N*_1_ and *N*_2_ are the number of solute molecules in the nucleus and cell, respectively. Therefore,

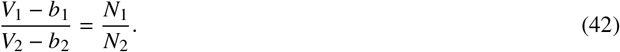

This equation is true regardless of the external concentration and so far we have not invoked any hypothesis on the *b_i_*. In particular, at the isotonic concentration at which 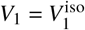 and 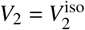, we have

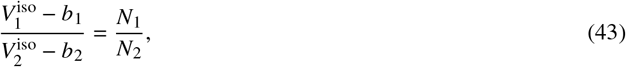

or equivalently

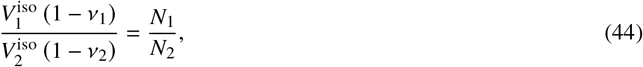

where 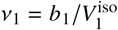 and 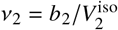 are dimensionless non-osmotic volumes.

It follows that the N/C ratio at the isotonic concentration, denoted by *NC*^iso^, satisfies

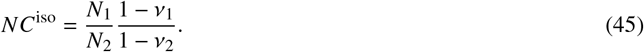

Next, we find a formula for the N/C ratio that holds for arbitrary external concentrations. We recall (42), which asserts that as a consequence of force-balance

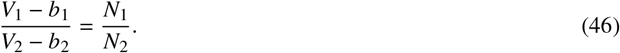

Multiplying through both sides by *V*_2_ – *b*_2_ yields

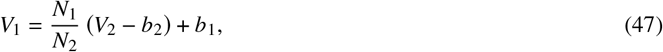

and dividing both sides by *V*_2_ results in

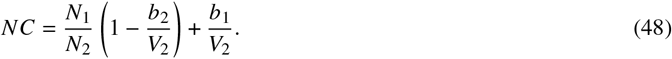

Straightforward algebra gives

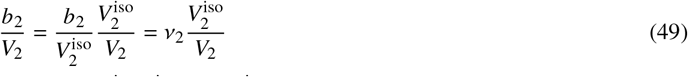

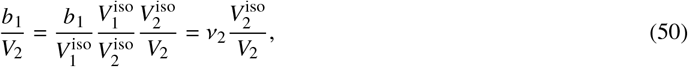

and plugging eqs. (49) and (50) into (48) yields

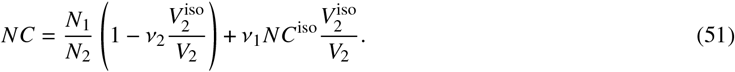

By (45), we have

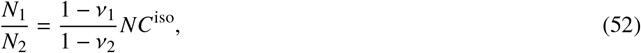

so that (51) may be rewritten as

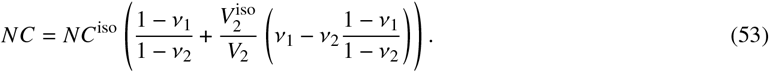

Adding and subtracting the term 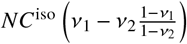 yields

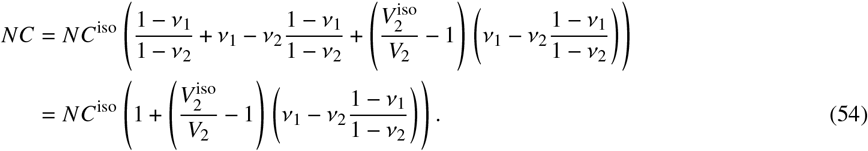

Next, we invoke the hypothesis 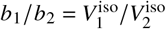, i.e. *v*_1_ = *v*_2_. It follows from (54) that if *v*_1_ = *v*_2_, in which case

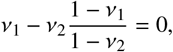

then *NC* = *NC*^iso^ independent of the external concentration and cell size *V*_2_.

On the other hand, if *v*_1_ ≠ *v*_2_, the N/C ratio will vary as 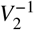 according to the formula given by (54).

